# Mutant Prion Protein Endoggresomes are Hubs for Local Axonal Organelle-Cytoskeletal Remodeling

**DOI:** 10.1101/2023.03.19.533383

**Authors:** Tai Chaiamarit, Adriaan Verhelle, Romain Chassefeyre, Nandini Shukla, Sammy Weiser Novak, Leonardo R. Andrade, Uri Manor, Sandra E. Encalada

**Affiliations:** Department of Molecular Medicine, The Scripps Research Institute, La Jolla, CA, 92037 USA; Dorris Neuroscience Center, The Scripps Research Institute, La Jolla, CA, 92037 USA; Neurodegeneration New Medicines Center, The Scripps Research Institute, La Jolla, 92037 CA, USA; Waitt Advanced Biophotonics Center, The Salk Institute for Biological Studies, La Jolla, 92037 CA, USA

## Abstract

Dystrophic axons comprising misfolded mutant prion protein (PrP) aggregates are a characteristic pathological feature in the prionopathies. These aggregates form inside endolysosomes -called endoggresomes-, within swellings that line up the length of axons of degenerating neurons. The pathways impaired by endoggresomes that result in failed axonal and consequently neuronal health, remain undefined. Here, we dissect the local subcellular impairments that occur within individual mutant PrP endoggresome swelling sites in axons. Quantitative high-resolution light and electron microscopy revealed the selective impairment of the acetylated vs tyrosinated microtubule cytoskeleton, while micro-domain image analysis of live organelle dynamics within swelling sites revealed deficits uniquely to the MT-based active transport system that translocates mitochondria and endosomes toward the synapse. Cytoskeletal and defective transport results in the retention of mitochondria, endosomes, and molecular motors at swelling sites, enhancing mitochondria-Rab7 late endosome contacts that induce mitochondrial fission via the activity of Rab7, and render mitochondria dysfunctional. Our findings point to mutant Pr Pendoggresome swelling sites as selective hubs of cytoskeletal deficits and organelle retention that drive the remodeling of organelles along axons. We propose that the dysfunction imparted locally within these axonal micro-domains spreads throughout the axon over time, leading to axonal dysfunction in prionopathies.

## INTRODUCTION

Compelling evidence indicates that axonal pathologies contribute to neurodegeneration in Alzheimer’s disease (AD), Parkinson’s disease (PD), Huntington’s disease (HD), amyotrophic lateral sclerosis (ALS), and prion diseases^1–4^. Specifically, accumulations of misfolded and aggregate-prone proteins along axons are often observed at swelling sites, and these have been linked to neuronal dysfunction via poisoning of axonal transport^5, 6^, by trapping of organelles, proteins, and cytoskeletal elements, or by damage to mitochondrial dynamics and health^7–9^.

Importantly, axonal deficits precede clinical manifestations and occur before neuronal cell body loss, strongly implicating axonal dysfunction as one of the earliest pathological events in neurodegeneration^3, 10, 11^. However, the subcellular mechanisms leading to axonal damage in the proteinopathies remain largely undefined.

Sporadic, infectious, and familial prion diseases (PrDs) are progressive and fatal neurodegenerative disorders commonly associated with neuronal loss, spongiform change, astrogliosis, and microglial activation^12^. Brain pathological lesions occur following conversion of the glycosylphosphatidylinositol (GPI)–anchored cellular/wild-type prion protein (PrP^C^/PrP^WT^) into an aggregate-prone misfolded PrP conformation (PrP^Sc^-scrapie)^13–15^. PrDs are characterized by the deposition of misfolded PrP aggregates throughout the central nervous system (CNS), including within dystrophic swellings in axons^16^, where human disease-associated mutant PrP has been shown to actively form neurotoxic aggregates within enlarged endolysosomes called endoggresomes^17^. In addition to PrP aggregates, axonal swellings also contain trapped mitochondria and enlarged membrane-bound endosomes^18^, and these are commonly observed in axons of cultured primary neurons expressing disease-associated mutant PrP, in the brains of rodents infected with scrapie or that harbor PrP mutations, and in post-mortem brains of patients with human prion disease^9, 17–21^. The buildup of endosomes and organelles in axons suggest a breakdown of the axonal cytoskeletal and trafficking systems, but how the genesis of these putative pathological lesions relate to the presence of misfolded protein aggregates at localized regions in axons, remains unexplored.

Within the axon, tightly regulated endolysosomal trafficking and cytoskeletal-based transport systems ensure the faithful distribution of membrane-bound endolysosomal proteins and organelles over long distances^17, 22^. The coordinated activities of the molecular motors kinesin and dynein drive the anterograde (toward synapse), and retrograde (toward cell body) movement, respectively, of pre-synaptic vesicles, mitochondria, and of endosomes along microtubule (MT) tracks that line up along the length of axons^23^. Regulation of axonal transport is achieved in part by MT post-translational modifications (PTMs) to tubulin, including acetylation and tyrosination/detyrosination, which impart MT track mechanical stability and also serve as signals for the recruitment of MT associated proteins (MAPs), molecular motors, and other regulatory signaling molecules^24–26^. An intact transport system is critical for the proper translocation and distribution of endosomes and mitochondria along axons, including to distal synapses where their close interactions regulate energy production and calcium/metabolite buffering^27^. Tight coordination of motor activity also drives mitochondrial fusion and fission dynamics, key events that ensure the maintenance of functional axonal mitochondria, especially during cellular stress^28^. Impairments in mitochondrial fusion/fission dynamics has been proposed to contribute to the pathophysiology of PrDs^29, 30^, yet it is unclear whether misfolded protein aggregation including of PrP, directly drives organelle-endosome accumulations and mitochondrial dysfunction in axons.

In an effort to understand the mechanisms of axonotoxicity in the familial prionopathies, here, we dissect the subcellular changes that occur within local axonal swelling sites that contain enlarged misfolded PrP endoggresomes in a neuronal model of familial prion disease. We show that a disease-associated PrP mutation harboring a 14-octapeptide repeat insertion (PrP^PG14^)^31^ forms enlarged endoggresomes at axonal swelling sites that induce dysfunction to cytoskeletal MT stability uniquely within these localized domains. A cascade of selective poisoning of motor-dependent transport of mitochondria, pre-synaptic vesicles and other endosomes ensues, uniquely affecting their movement toward the synapse. Axonal swellings enriched with enlarged PrP endoggresome also actively trap molecular motors, mitochondria, and endosomes, enhancing endosome-mitochondrial contacts that directly drive local endosome-mediated mitochondrial fission. These observations suggest that axonal swelling sites containing misfolded PrP endoggresomes serve as axonotoxicity hubs that unleash cascades of selective cytoskeletal and motor destabilization, trigger disruptions in organelle movement, and are magnets for endosome-organelle assemblages that promote mitochondrial remodeling towards fission resulting in failed mitochondrial health. Importantly, the disruption of cytoskeletal and endosomal pathways within local axonal domains occur at early stages following initial expression of PrP mutations. Thus, the cascade of localized eddies of impairments that occur within swelling/aggregate sites in axons explain how axons are uniquely vulnerable to the formation of aggregates and how these aggregates trigger axonal demise. Moreover, these studies uncover local cytoskeletal and organelle-organelle dysfunction pathways as potential early targets of intervention for the development of therapies aimed at reversing these local axonal deficits in the PrDs.

## RESULTS

### Axonal transport is differentially impaired in axons with PrP^PG14^ endoggresomes

Our previous work showed that expression of PrP^PG14^-mCherry (mCh) or -EGFP in hippocampal murine neurons resulted in the formation of large misfolded PrP^PG14^ aggregates inside endolysosomes—called endoggresomes—at localized swelling sites along axons^17^ (**Supplementary Fig. 1a)**. PrP^PG14^-EGFP endoggresomes were observed in axons 12 hours post-transfection, and endoggresome densities increased over 2 days of expression, while axons expressing PrP^WT^-EGFP contained primarily small vesicles that were highly motile (**Supplementary Fig. 1b**). Densities of PrP^PG14^-EGFP and PrP^PG14^-mCh endoggresomes were comparable 2 days post-expression^17^. To investigate whether expression of PrP^PG14^-mCh resulted in impaired axonal function, we characterized the axonal transport of three cargoes that actively mobilize in mammalian axons and that are required for proper axonal function—mitochondria, pre-synaptic vesicles, and PrP vesicles^32–34^. Single particle transport of these cargoes was imaged at high spatial and temporal resolution in axons of neurons co-transfected with PrP^PG14^-mCh or PrP^WT^-mCh and either mito-EGFP (containing the mitochondrial targeting sequence of Cytochrome c Oxidase subunit VIII fused to EGFP) or SYP-YFP (Synaptophysin fused to YFP; **Methods**). Time-lapse videos were analyzed using a custom-made single particle tracking analysis software to generate trajectory datasets^35^. At two days post-transfection, the densities of motile mito-EGFP particles were significantly reduced in PrP^PG14^-mCh axons compared to PrP^WT^-mCh controls (**Fig. 1a,b**), suggesting defective mitochondrial translocation into the axon. To further test whether the mitochondrial bidirectional movement was impaired, analysis of segmental trajectories 2 days post-expression revealed that mito-EGFP spent less time moving in the anterograde direction and more time pausing in PrP^PG14^-mCh endoggresome-rich axons compared to PrP^WT^-mCh aggregate-free axons (**Fig. 1c**). These mito-EGFP particles also undertook shorter anterograde run lengths (**Fig. 1d**) and had reduced anterograde instantaneous velocities (**Fig. 1e**). Notably, retrograde transport dynamics of mito-EGFP particles remained unchanged in PrP^PG14^-mCh endoggresome-rich axons compared to PrP^WT^-mCh axons (**Fig. 1c-e**). These data show differential deficits in the movement of mitochondria uniquely towards the synapse. Notably, despite gradual accumulation of PrP^PG14^-mCh endoggresomes 12 hours after initial PrP^PG14^-mCh expression (**Supplementary Fig. 1b**), mito-EGFP transport was largely unaltered at this time point and thus was comparable to that in PrP^WT^-mCh control axons (**Fig. 1a-c**), with an exception in the run lengths (**Fig. 1d**). These data indicate that mitochondrial transport impairment lags behind but progresses with time alongside PrP^PG14^ aggregation.

**Fig. 1:**
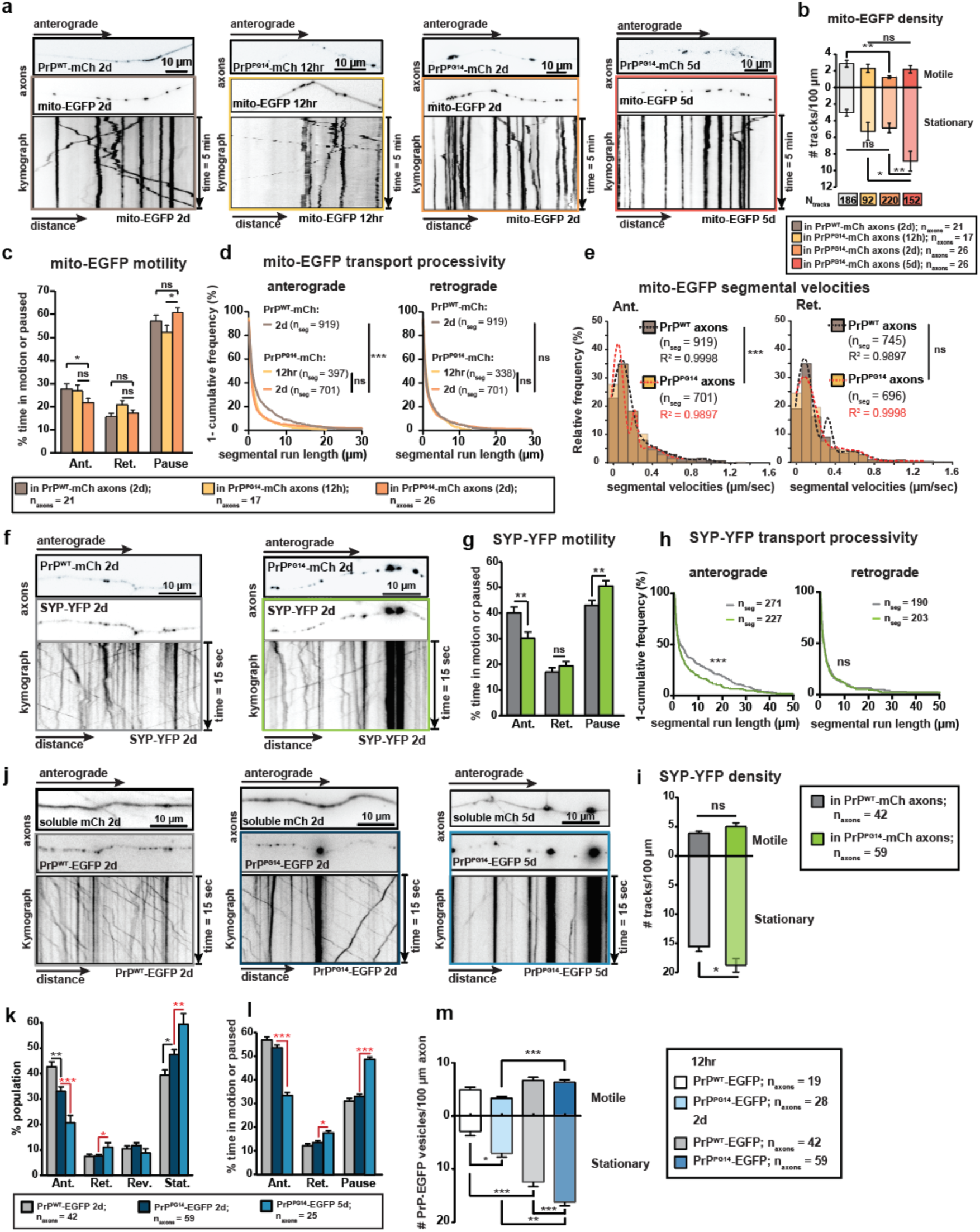
Axonal transport is progressively impaired in PrP^PG14^-mCh axons. **a** First-frame images of axons co-transfected with mito-EGFP and PrP^WT^-mCh or PrP^PG14^-mCh for 12 hours or 2 days (top two panels) and corresponding kymographs (bottom panel) generated from time-lapse movies (5 minutes, 300 frames, 1 frame/sec). **b** Densities of motile (top y-axis) or stationary (bottom y-axis) mito-EGFP. **c** Percent time mito-EGFP spends in anterograde (Ant.) or retrograde (Ret.) motion or paused. **d** Segmental run lengths of mito-EGFP plotted as 1-cumulative frequency (%). **e** Segmental velocities of mito-EGFP plotted as relative frequency histogram (%) and fitted with multi-peaks Gaussian functions. R^2^ values indicate the degree of Gaussian fits. **f** First-frame images of axons co-transfected with SYP-YFP and PrP^WT^-mCh or PrP^PG14^-mCh for 2 days (top two panels) and corresponding kymographs (bottom panel) generated from time-lapse movies (15 seconds, 150 frames, 10 frame/sec). **g** Percent time SYP-YFP spends in anterograde (Ant.), or retrograde (Ret.) motion, or paused. **h** Segmental run lengths of SYP-YFP plotted as 1-cumulative frequency (%). **i** Densities of motile (top y-axis) or stationary (bottom y-axis) SYP-YFP. **j** First-frame images of axons co-transfected with soluble mCh and PrP^WT^-EGFP or PrP^PG14^-EGFP for 2 or 5 days (top two panels) and corresponding kymographs (bottom panel) generated from time-lapse movies (15 seconds, 150 frames, 10 frame/sec). **k** Percent population with anterograde (Ant.), retrograde (Ret.), reversal (Rev.), or stationary (Stat.) tracks. **l** Percent time PrP^WT^-EGFP or PrP^PG14^-EGFP spend in anterograde (Ant.), or retrograde (Ret.) motion, or paused. **m** PrP^WT^-EGFP or PrP^PG14^-EGFP densities of motile (top y-axis) or stationary (bottom y-axis) vesicles. All bar graphs are shown as mean ± SEM. ***p<0.001, **p<0.01, *p<0.05, ns = not significant. **b**, **c, k, l, m** Student’s t-test with Tukey’s multiple comparison tests; **g, i** Student’s t-test; **d**, **e**, and **h** Kolmogorov-Smirnov (K-S) test.

Mitochondria utilize kinesin-1 (KIF5 or kinesin heavy chain [KHC]) to translocate anterogradely to the axonal terminus^27^. However, reduction of a neuronally-enriched kinesin-3 (KIF1A) using a validated shRNA resulted in significant impairments to mitochondrial segmental run lengths and velocities as compared to those in D11 scrambled shRNA controls (**Supplementary Fig. 2a-c**), indicating that KIF1A is also an important driver of mitochondria movement in axons of hippocampal neurons. As the overall anterograde transport of mito-EGFP was defective upon expression of PrP^PG14^-mCh (**Figure 1a-e**), this raises the possibility that both kinesin-1- and kinesin-3-driven transport dynamics are impaired in PrP^PG14^-mCh-expressing axons.

We next analyzed the movement of another KIFA cargo, pre-synaptic vesicles carrying SYP-YFP, an integral membrane protein that is trafficked to the pre-synapse and is important for vesicle fusion, synaptic vesicle endocytosis, and recycling^36, 37^. Transport dynamics of SYP-YFP vesicles 2 days post-transfection were analyzed using time-lapse quantitative fluorescence microscopy at high spatial and temporal resolution (**Fig. 1f; Methods**). In comparison to axons expression PrP^WT^-mCh, the percent time that SYP-YFP vesicles spent in anterograde motion in PrP^PG14^-mCh axons was significantly decreased, while the time spent paused was increased (**Fig. 1g**). SYP-YFP vesicles also had significantly shorter segmental run lengths in PrP^PG14^-mCh axons compared to PrP^WT^-mCh control axons (**Fig. 1h**). As with mitochondrial transport, retrograde transport of SYP-YFP vesicles was also unaffected in the endoggresome-rich PrP^PG14^-mCh axons (**Fig. 1g-h**), but a significantly higher proportion of SYP-YFP vesicles remain stationary in PrP^PG14^-mCh axons, compared to PrP^WT^-mCh axons (**Fig. 1i**), suggesting the overall retention of SYP-YFP transport towards the distal synapse.

In addition to forming PrP^PG14^ endoggresomes, axons of neurons expressing PrP^PG14^-mCh contain motile Golgi-derived PrP^PG14^-mCh vesicles that fuse in axons during active transport toward the synapse to form PrP^PG14^ endoggresomes (**Fig. 1j**; see **Methods** for definition of PrP^PG14^ vesicles versus endoggresomes)^17^. Quantitative single particle tracking analysis showed that 2 days post-expression, the percent anterograde-moving PrP^PG14^-mCh vesicles in axons significantly decreased, while the percent of retrograde and stationary populations increased (**Fig. 1j,k**). These differences were amplified 5 days post-expression. At the 5-day time point, the time that PrP^PG14^-mCh vesicles spent in motion in the anterograde direction was also significantly decreased, while more time was spent by PrP^PG14^-mCh vesicles moving in the retrograde direction or paused (**Fig. 1l,m**). Altogether, starting at 12 hours post-transfection of PrP^PG14^-mCh, gradual increases in transport deficits of PrP^PG14^-mCh vesicles were observed, while mito-EGFP and SYP-YFP vesicle transport was impaired at 2 days post-transfection (**Fig. 1a-h**). These observations show that early transport defects exhibited by PrP^PG14^-mCh vesicles concur with growing PrP^PG14^-mCh endoggresome densities in axons at that time point (**Supplementary Fig. 1b**), suggesting that the movement of PrP^PG14^-mCh vesicles that are being removed from the motile vesicle pool to fuse and form enlarged endoggresomes^17^, is progressively affected at very early stages. Moreover, the differential impairments uniquely to the anterograde transport of mitochondria and pre-synaptic suggest that anterograde transport is more susceptible to the presence of PrP^PG14^ aggregates in axons, and these observations raise the possibility of differential impairments to the anterograde transport machinery.

### Local traffic jams at axonal PrP^PG14^ aggregate sites revealed by micro-movement analysis

As transport of mitochondria was impaired resulting in increased number of stalled mitochondria in axons of neurons expressing PrP^PG14^ (**Fig. 1a-c**), we next asked whether PrP^PG14^ endoggresomes directly hindered the movement of mitochondria and retained these organelles within these local swelling sites. We used single-particle tracking to image and quantitate the behavior of individual mitochondria within micro-domain regions at PrP^PG14^ endoggresome/aggregates sites (herein referred as ‘PrP^PG14^ aggregate sites or AGG) vs non-aggregate regions (NON-AGG) (**Fig. 2a**). AGG and NON-AGG regions within the same axon were defined within 10-µm wide analysis windows (**Fig. 2a, d**). At each window, we quantified percentages of mito-EGFP that (1) passed through AGG sites without interruptions, (2) paused/slowed down at the AGG sites, and (3) stalled at the AGG sites, compared to those in control NON-AGG sites of axons of PrP^PG14^-mCh-expressing neurons in the anterograde versus retrograde directions (**Fig. 2a-c**). Our data indicate that while ∼50% of the mito-EGFP population stalled along axons (**Fig. 2c**), from those that retained motility, a significantly lesser proportion moved through AGG sites without interruption compared to NON-AGG sites (**Fig. 2b**). Moreover, a significantly greater proportion of moving mitochondria paused at- or slowed down as they moved in the anterograde direction through AGG vs NON-AGG sites (**Fig. 2b**). These data suggest that PrP^PG14^ AGG sites are magnets for mitochondria, to promote the arrest or the slowing down of anterograde-moving mitochondria. To further characterize the degree of mitochondrial arrest at PrP^PG14^ AGG sites, mitochondrial micro-movement dynamics were quantified within 10 µm-wide windows surrounding individual AGG or NON-AGG sites (**Fig. 2d**). We measured the mean displacement of individual mito-EGFP from their centroid position over a x min period of live imaging, within 10 µm AGG vs NON-AGG windows (**Fig. 2d**). Compared to NON-AGG regions, single particle tracking revealed that mito-EGFP displacement was largely within the 10 µm window (<10 µm), thus mitochondria did not move as far (**Fig. 2e**). Analysis of segmental velocities within the 10 µm windows revealed that mito-EGFP at AGG sites moved significantly slower (<0.1 µm/s) compared to mitochondria at NON-AGG sites (**Fig. 2f**). Altogether, these data suggest that the mitochondrial motility impairments represent local ‘traffic jams’ at PrP^PG14^ endoggresome AGG sites and suggest that the global deficits in axonal transport of mitochondrial along axons result from micro-transport disruptions that start at AGG sites where PrP^PG14^ endoggresomes themselves are stalled, thus implicating PrP^PG14^ endoggresomes directly in the initiation of these local transport deficits.

**Fig. 2:**
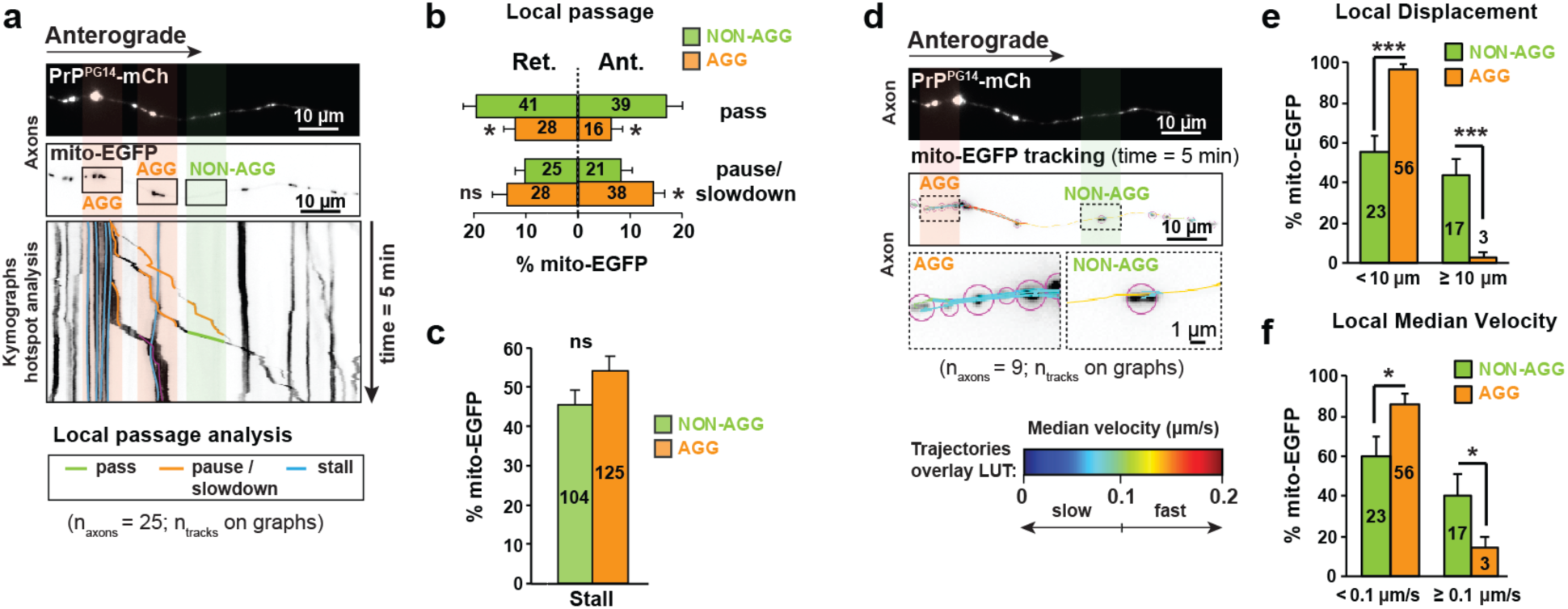
Local axonal transport disruption of mito-EGFP micro-movements within PrP^PG14^-mCh aggregate sites. **a** First-frame images of an axon co-transfected with PrP^PG14^-mCh (top) and mito-EGFP (middle) and corresponding kymograph from a time-lapse movie (5 minutes, 300 frames, 1 frame/sec) of mito-EGFP transport with color-coded track assignment overlay (green = pass, orange = pause/slowdown, or blue = stall) at AGG and NON-AGG sites. **b** Percent number of mito-EGFP that pass or pause/slowdown in anterograde (Ant.; right x-axis) vs retrograde (Ret.; left x-axis) directions at AGG vs NON-AGG sites. **c** Percent number of mito-EGFP that are stalled at AGG vs NON-AGG sites. **d** First-frame images of an axon co-transfected with PrP^PG14^-mCh (top) and mito-EGFP (bottom) from time-lapse movies (5 minutes, 300 frames, 1 frame/sec). Dotted rectangles outline 10 µm-wide PrP^PG14^ aggregate sites (AGG) or non-aggregate control sites (NON-AGG). Rainbow LUT pseudo color-coded overlays indicate median velocities ranges of tracks at AGG and NON-AGG regions (insets). **e** Percent number of mito-EGFP trajectories at AGG vs NON-AGG sites with track displacements of < 10 µm or ≥ 10 µm during a 5-minute acquisition period. **f** Percent number of mito-EGFP at AGG vs NON-AGG sites with median velocity of < 0.1 µm/s or ≥ 0.1 µm/s during the 5-minute acquisition. All bar graphs are shown as mean ± SEM. ***p<0.001, **p<0.01, *p<0.05, ns = not significant. **b**, **c**, **e**, **f** Student’s t-test. Number of tracks are shown inside bar graphs. All analyses were done on axons after 2 days of PrP^PG14^-mCh transfection.

### Local changes to the actin and MT cytoskeleton drive at PrP^PG1^^4^ aggregate sites

The cylindrical shape of the axon is supported by a tightly organized and regularly spaced actin-spectrin lattice localized below the plasma membrane, and by a system of MTs that line up the length of the axon^38, 39^. An intact actin and MT cytoskeletal organization is a key determinants of axonal transport dynamics^39, 40^. To investigate the mechanism(s) of mitochondrial arrest and transport impairments at AGG sites (**Fig. 1**), we characterized the actin and MT cytoskeletal integrity at localized axonal PrP^PG14^ AGG swelling sites. The ultrastructure of actin and MT tracks inside axons was visualized using correlative light and scanning electron microscopy (CLEM) of cultured neurons grown on gridded coverslips. These cells were transfected with PrP^PG14^-mCh for 2 days, subjected to a membrane-extraction (ME) protocol to remove lipids including the plasma membrane, and imaged using SEM. In this approach, the position of individual PrP^PG14^-mCh endoggresomes was mapped within axons using fluorescence imaging, followed by treatment of live neurons with ME detergent buffers to remove lipid membranes, but also containing taxol to stabilize MTs and phalloidin to stabilize actin (**Fig. 3a**)^41^. Scanning electron micrographs revealed ring-like accumulations of actin surrounding, or localized with PrP^PG14^-mCh aggregates (**Fig. 3a, b; Supplementary Fig. 3a**). Similar actin ring-like structures were observed using direct stochastic optical reconstruction microscopy (dSTORM) imaging of Phalloidin-AF647 at PrP^PG14^-mCh AGG swellings (**Fig. 3b**), suggesting the active recruitment of actin to PrP^PG14^-mCh aggregate AGG sites.

**Fig. 3:**
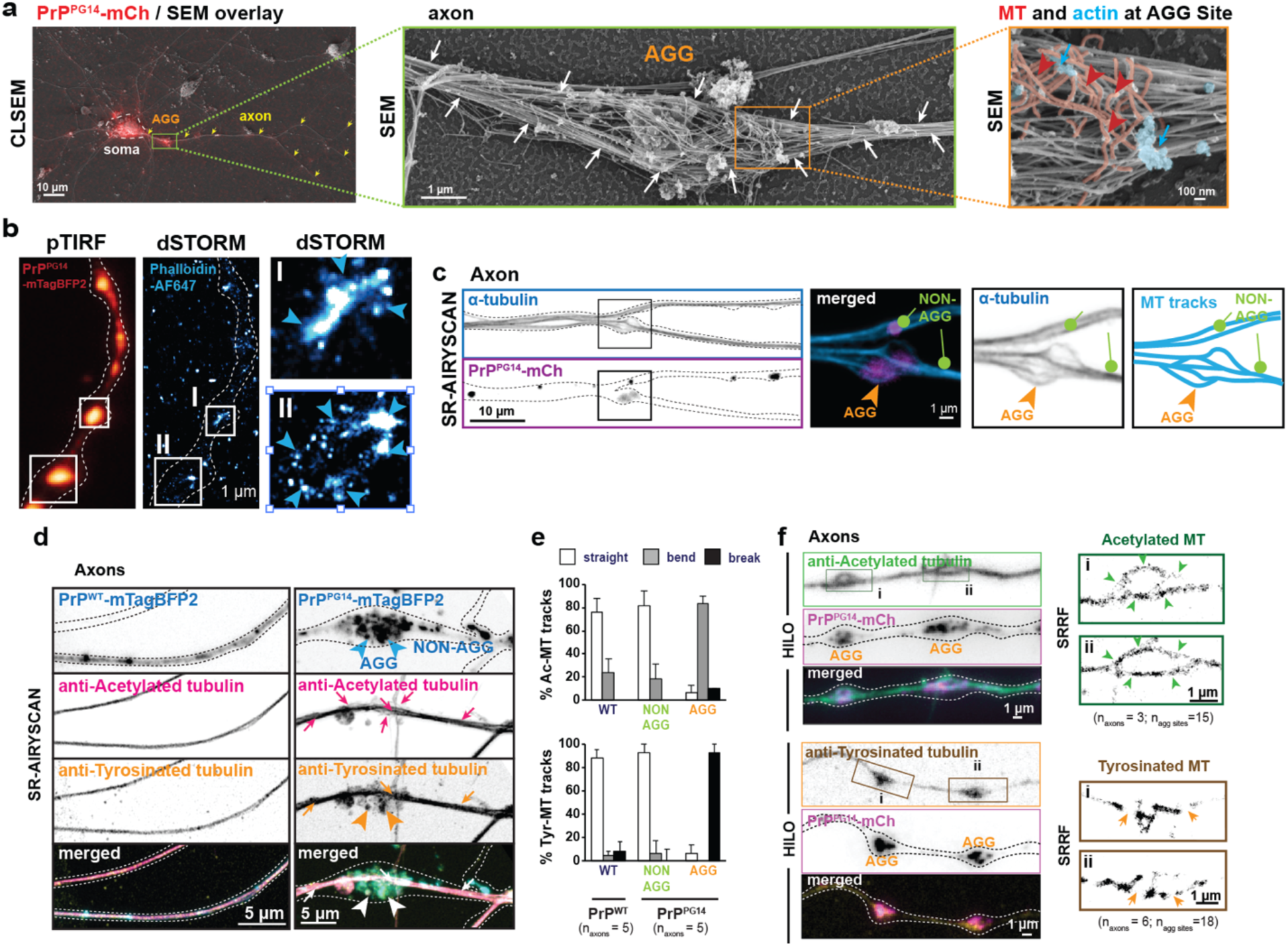
Actin accumulation and selective impairment to MT track PTMs at PrP^PG14^ aggregate sites. **a** Membrane-extracted (ME) correlative light and scanning electron microscopy (CLSEM) image of a neuron transfected with PrP^PG14^-mCh (red). Arrowheads indicate axons. Green Inset shown on middle panel. White arrows point to MT tracks. Orange inset shown on right panel. Blue arrows point to actin. Red arrowheads point to disrupted MT tracks. **b** Images of an axon from a neuron transfected with PrP^PG14^-mTagBFP2 (red) imaged using highly-inclined and laminated optical sheet (HILO) microscopy, stained with Phalloidin-AF647 and imaged with direct STORM (dSTORM). Insets I-II of axonal swellings shown on the right. Arrowheads point to actin accumulation around the axonal swellings. **c** Super-resolution (SR) Airyscan confocal microscopy maximum projection images of PrP^PG14^-mCh, immunostained with alpha tubulin antibody. Round Arrows indicate MT tracks at NON-AGG sites; Arrowheads indicate MT at AGG site. **d** SR-Airyscan images of PrP^WT^- or PrP^PG14^-mTagBFP2 axons, co-immunostained with acetylated (Ac) and tyrosinated (Tyr) tubulin antibodies. Arrows indicate intact/straight tracks; Arrowheads indicate broken MT. **e** Percent Ac- or Tyr-MT track architecture (straight, bend, or break) at WT (in PrP^WT^-mTagBFP2 axons) and NON-AGG or AGG sites (in PrP^PG14^-mTagBFP2 axons). **e** HILO microscopy images of PrP^PG14^-mCh axons immunostained against either acetylated and tyrosinated tubulin, and corresponding Super-resolution radial fluctuation (SRRF)-processed images at the AGG sites (i-ii; insets). Insets: Arrowheads point to MT tracks bending; Arrows point to MT tracks breaking. See also figure S3.

Axonal swelling sites also contained bending MTs that followed the contour of swellings at PrP^PG14^-mCh AGG sites, as well as broken MTs that curled up/broke at AGG sites, while NON-AGG neighboring regions had straight MT bundles (**Fig. 3a, Supplementary Fig. 3a, b**). These observations were confirmed using 3D super-resolution (SR)-Airyscan imaging of α-tubulin that showed curved MT tracks around at AGG sites in contrast to straight MTs observed at NON-AGG regions (**Fig. 3c, Supplementary Fig. 3b**). MT stabilization is imparted by post-translational modifications (PTMs) to tubulin^42^. α-tubulin acetylation is a common PTM that confers MT stability and protects MTs from breakage under mechanical stress^43^. Acetylated MTs are selectively preferred by kinesin-1 for motility^44–46^. On the other hand, tyrosinated (unmodified) MT tracks that are less stable and prone to breakage upon stress, and have existing tyrosine tails on tubulin often associated with kinesin-3, a motor reported to prefer tyrosinated MTs for motility usually though interactions with other MAPs^47–49^. Our observations of bending and breaking MTs prompted us to ask whether MT tracks with different PTMs were selectively impaired within PrP^PG1^^4^ endoggresome AGG sites. PrP^PG1^^4^-mCh axons were immunostained with validated tubulin antibodies that recognize acetylated vs tyrosinated MTs. 3D SR-Airyscan, and a highly inclined and laminated optical sheet (HILO) with super-resolution radial fluctuation (SRRF) processing^50^ revealed that most acetylated MT tracks bent or remained undisrupted while tyrosinated MT tracks broke at AGG sites, compared to NON-AGG sites (**Fig. 3d-f, Supplementary Fig. 3c**). These results are consistent with previous reports showing mechanically stable and breakage-resistance acetylated MT tracks^43^. Overall, these results show that actin and MTs are remodeled at the axonal swelling sites containing PrP^PG14^ endoggresomes. Furthermore, MT tracks with different PTMs are differentially impaired at the AGG sites. Because kinesin-1 and kinesin-3 recognize MT tracks with distinct PTMs, differential disruption of MTs at AGG sties could impinge on axonal transport dynamics of cargoes that depend on kinesin-1 and kinesin-3 for motility in axons.

### Selective retention of kinesin-1 and kinesin-3 at PrP^PG1^^4^ aggregate sites

The observed local and selective disruption of mitochondrial transport (**Fig. 2**), as well as the impaired MTs stability within PrP^PG14^ endoggresomes AGG sites (**Fig.3**), suggest defective interactions between molecular motors and cargoes at those sites. We first tested whether kinesin-1 (KHC - KIF5) and its associated adaptor subunit kinesin light chain 1 (KLC1), and kinesin-3 (KIF1A), two neuronally-enriched anterograde motors that bind to cargoes via adaptors and become activated in order to drive processive transport^51^, were aberrantly retained within AGG sites. We used immunofluorescence and validated antibodies to quantitate fluorescence intensity distributions of endogenous KHC, KLC1, and KIF1A at the AGG sites, as a measure of the relative amount of KHC and KIF1A motors present at those sites (**Fig. 4a**). These intensity frequency distributions were compared to (1) those at control NON-AGG sites in axons of PrP^PG14^-mCh-expressing neurons, and to (2) sites in axons of control PrP^WT^-mCh-expressing neurons (‘WT sites’) (**Fig. 4a**). As shown previously, KHC, KLC1, and KIF1A staining in axons of neurons expressing PrP^WT^-mCh was punctate^34, 52^, indicating that these proteins were localized to vesicular structures (**Fig. 4a**). However, intensities of KHC, KLC1, and KIF1A motors were higher at AGG sites, indicating the accumulation of motors within AGG swellings containing PrP^PG14^-mCh endoggresomes (**Fig. 4a**). Quantitation of fluorescence intensity distributions confirmed these observations: comparison of fluorescence intensity distribution graphs between AGG and NON-AGG sites showed significantly wider and right-skewed distributions toward higher intensities for KHC, KLC1, and KIF1A at AGG sites compared to WT sites (**Fig. 4b**). Notably, the KLC1 intensity distribution graph at NON-AGG sites was significantly lower compared to those at WT sites, suggesting that KLC1 was depleted from cargo moving through NON-AGG areas.

**Fig. 4:**
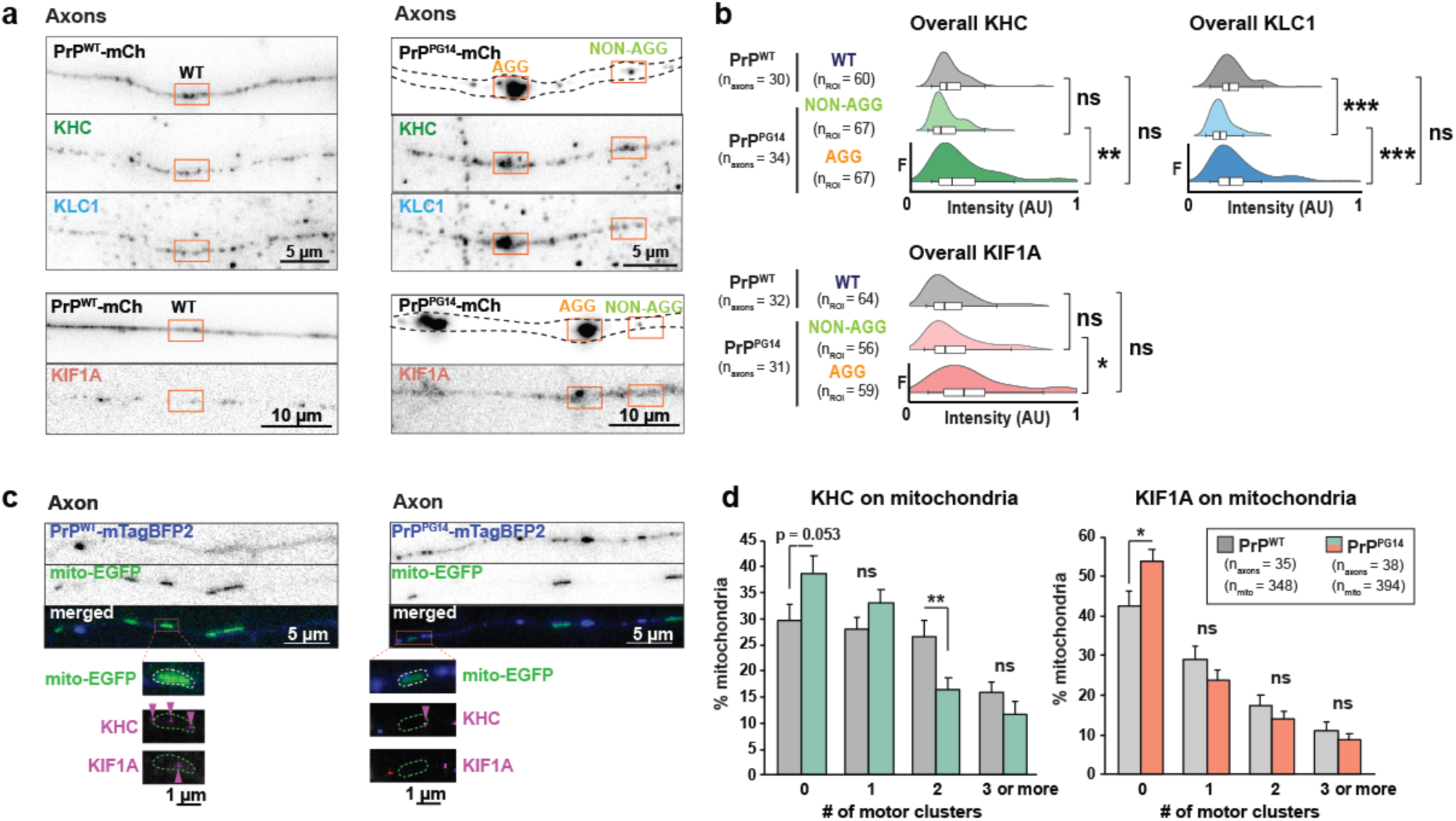
Kinesin-1 and kinesin-3 accumulation at PrP^PG14^ aggregate sites and depletion from mitochondria. **a** Images of axons expressing PrP^WT^-mCh or PrP^PG14^-mCh, immunostained with KHC and KLC1 or with KIF1A antibodies. Orange boxes show AGG, NON-AGG, and WT ROIs. **b** Normalized intensity distribution frequencies (F) of KHC, KLC1, and KIF1A signal at AGG, NON-AGG, and WT sites plotted as box plots with kernel density estimates from histogram. **c** Representative inverted images of axons expressing PrP^WT^-mTagBFP2 or PrP^PG14^-mTagBFP2 and mito-EGFP, and immunostained with KHC and KIF1A antibodies. Insets (below) shown as overlay of sub-pixel localization detection (magenta spots; arrowheads). **d** Quantitation of percent mitochondria (mean ± SEM) colocalizing with 0, 1, 2, and ≥ 3 KHC or KIF1A motor clusters. ***p<0.001, **p<0.01, *p<0.05, ns = not significant; **b** Kruskal-Wallis test, **d** Student’s t-test.

Next, to test whether motor-cargo association was disrupted as a result of motor sequestration at the AGG sites, we analyzed kinesin-1 and kinesin-3 motor composition on individual mitochondria that mobilized through PrP^PG14^ endoggresome AGG sites, and compared this to the composition of motors associated with mitochondria in PrP^WT^ aggregate-free control axons. We used a sub-pixel localization algorithm^34, 53^ to detect and compare the degree of colocalization between KHC and KIF1A motor subunits on mitochondria labeled with mito-EGFP) (**Fig. 4c**). In this analysis, diffraction-limited puncta representing ‘motor clusters’ were analyzed containing one or more kinesin molecules. Our data show that mito-EGFP that had moved through at least one PrP^PG14^ endoggresome AGG site contained significantly less KHC and KIF1A motor clusters, compared to mito-EGFP in PrP^WT^ control axons (**Fig. 4d**). As mitochondria analyzed here had moved through at least than one AGG site, these observations suggest that kinesin-1 and kinesin-3 are depleted away from mitochondria as they passage through PrP^PG14^ AGG swellings, resulting in kinesin-1 and kinesin-3 accumulation at these sites.

### Retention of kinesin-1 at PrP^PG1^^4^ aggregate sites requires its cargo-binding domain

As the cargo-binding KLC1 subunit of kinesin-1 was depleted from mitochondrial cargo that passed through PrP^PG14^ endoggresome AGG sites (**Fig. 4b**), to determine the mechanistic basis of motor sequestration at PrP^PG14^ AGG sites, we tested whether a well characterized truncated kinesin-1 (KIF5C[1-560]) consisting of the motor, neck linker, neck coil, and first stalk domains, but that lacks the cargo binding domain and thus cannot associate with cargoes^54^, was retained at PrP^PG14^ AGG sites. Neurons were co-transfected with PrP^PG14^-mTagBFP2 (mTag blue fluorescence protein 2) and full-length (FL) KIF5C-2xmCh plus its adaptor subunit KLC1-TAP, or with a KIF5C(1-560)-2xmCh plus KLC1-TAP (**Fig. 5**). We found that KLC1-TAP overexpression was sufficient to recruit the FL KIF5C-2xmCh to PrP^PG14^ endoggresomes (**Fig. 5a**). These data are consistent with immunofluorescence image analysis of endogenous KIF5 (KHC) and KLC1 showing accumulation of these motors at PrP^PG14^ endoggresome AGG sites (**Fig. 4a,b**). In contrast, truncated KIF5C(1-560)-2xmCh did not accumulate at PrP^PG14^ AGG sites throughout axons, despite KLC1 overexpression (**Fig. 5b**). Instead, KIF5C(1-560)-2xmCh efficiently moved passed all AGG sites through the axon to accumulate at axonal termini **(Fig. 5c**) in a similar manner as previous observed in healthy cultured neurons^54^. Uninterrupted movement of KIF5C(1-560) is consistent with our observation that kinesin-1 prefers acetylated MTs, and these remained mostly intact and curved at the AGG sites (**Fig. 3d-f**), thus providing the tracks for KIF5C(1-560) transport toward the axon terminus. Next, although the movement of KIF5C(1-560) was not blocked, we tested whether motility dynamics of KIF5C(1-560) were impaired as it moved through PrP^PG14^ AGG sites before reaching the axon terminals.

**Fig. 5:**
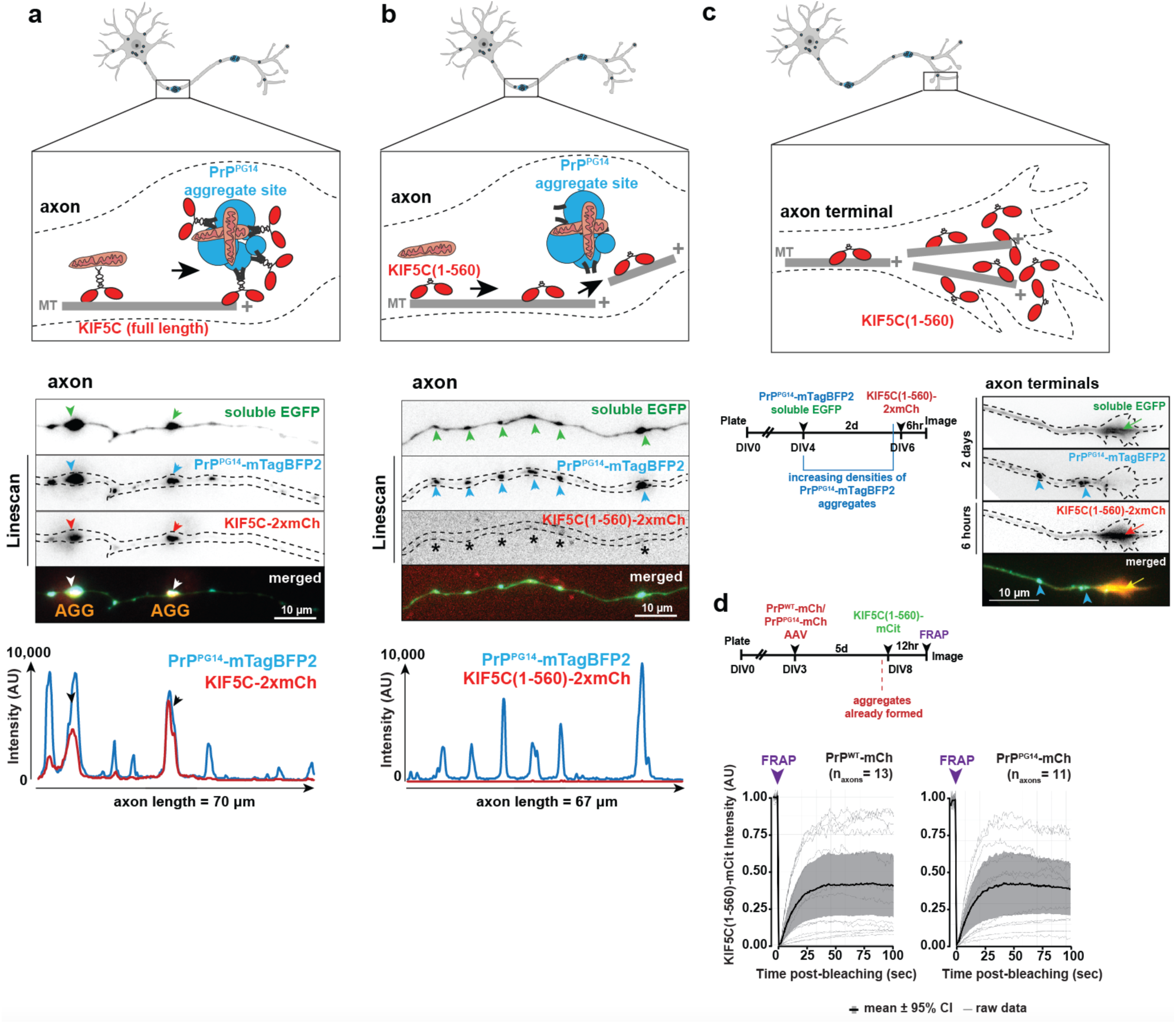
Retention of kinesin-1 (KIF5C) at PrP^PG14^ AGG sites requires its cargo-binding domain. **a** Schematic (top) and representative inverted images of axons (middle) of neurons expressing soluble EGFP, PrP^PG14^-mTagBFP2, and KIF5C-2xmCh for 2 days, and corresponding line scan intensity profiles (bottom) (n_axons_ = 9). **b** Schematic (top) and representative inverted images of axons (middle) from neurons expressing soluble EGFP, PrP^PG14^-mTagBFP2, and KIF5C(1-560)-2xmCh for 2 days, and corresponding line scan intensity profiles (bottom) (n_axons_ = 13). **c** Schematic (top), timeline of sequential transfection of EGFP and PrP^PG14^-mTagBFP2 for 2 days and KIF5C(1-560)-2xmCh for 6 hours (bottom left), and images at the axon terminus (bottom right). Arrowheads point to endoggresomes. Arrows point to EGFP and KIF5C(1-560)-2xmCh accumulation at the axonal growth cone. **d** Timeline of FRAP assay (top) of axon terminals induced with PrP^WT^- or PrP^PG14^-mCh AAV for 5 days, sequentially transfected with KIF5C(1-560)-mCit for 12 hours prior to live imaging. (bottom) FRAP profile of KIF5C(1-560)-mCit.

Fluorescence recovery after photobleaching (FRAP) analysis of KIF5C(1-560)-mCitrine (mCit) at the distal tips of axons of neurons expressing PrP^PG14^-mCh vs. PrP^WT^-mCh using adeno associated viruses (AAVs) 5 days post-transduction, a time point previously shown to form prominent aggregates^17^, showed no significant difference in the FRAP recovery of KIF5C(1-560)-mCit in PrP^PG14^ endoggresome-rich axons compared to control PrP^WT^ axons devoid of endoggresomes (**Fig. 5c**), further suggesting that PrP^PG14^ endoggresomes do not block the transport of KIF5C(1-560) to the axonal termini. These data show that the cargo-binding domain of KIF5C is sufficient and required for retention of cargo-motor complexes at PrP^PG14^ AGG swelling sites in axons. These observations suggest that there is selectivity in this retention, as motors without cargo-binding domains, i.e., without cargo, including soluble motors are not retained within AGG sites, thus indicating that only intact cargo-motor complexes are sequestered at PrP^PG14^ AGG sites.

### Enhanced mitochondrial fission at PrP^PG1^^4^ aggregate sites requires Rab7 GTPase activity

In addition to mitochondrial accumulation (**Fig. 2**), time-lapse live imaging using high-resolution microscopy revealed enhanced mitochondrial fission at PrP^PG14^ AGG sites (**Fig. 6a**, **Supplementary Video 1)**. To characterize mitochondrial fission, we first quantitated mitochondrial size at PrP^PG14^ AGG sites in axons of neurons expressing PrP^PG14^-mCh for 5 days (**Fig. 6a**). We observed smaller mitochondria within PrP^PG14^ AGG sites (**Fig. 6b**), and while many smaller mitochondrial fragments were retained within AGG sites, some also moved outside of the AGG site in the anterograde direction (**Fig. 6a**). Moreover, the size of mitochondria trapped within PrP^PG14^ AGG sites decreased as the number of PrP^PG14^ endoggresomes increased from 12 hours to 5 days following initial expression of PrP^PG14^-mCh (**Fig. 6c,d, Supplementary Fig. 1b**). These data suggest a link between the presence of PrP^PG14^ endoggresomes and the progressive imbalance of mitochondria fission dynamics at AGG sites.

**Fig. 6:**
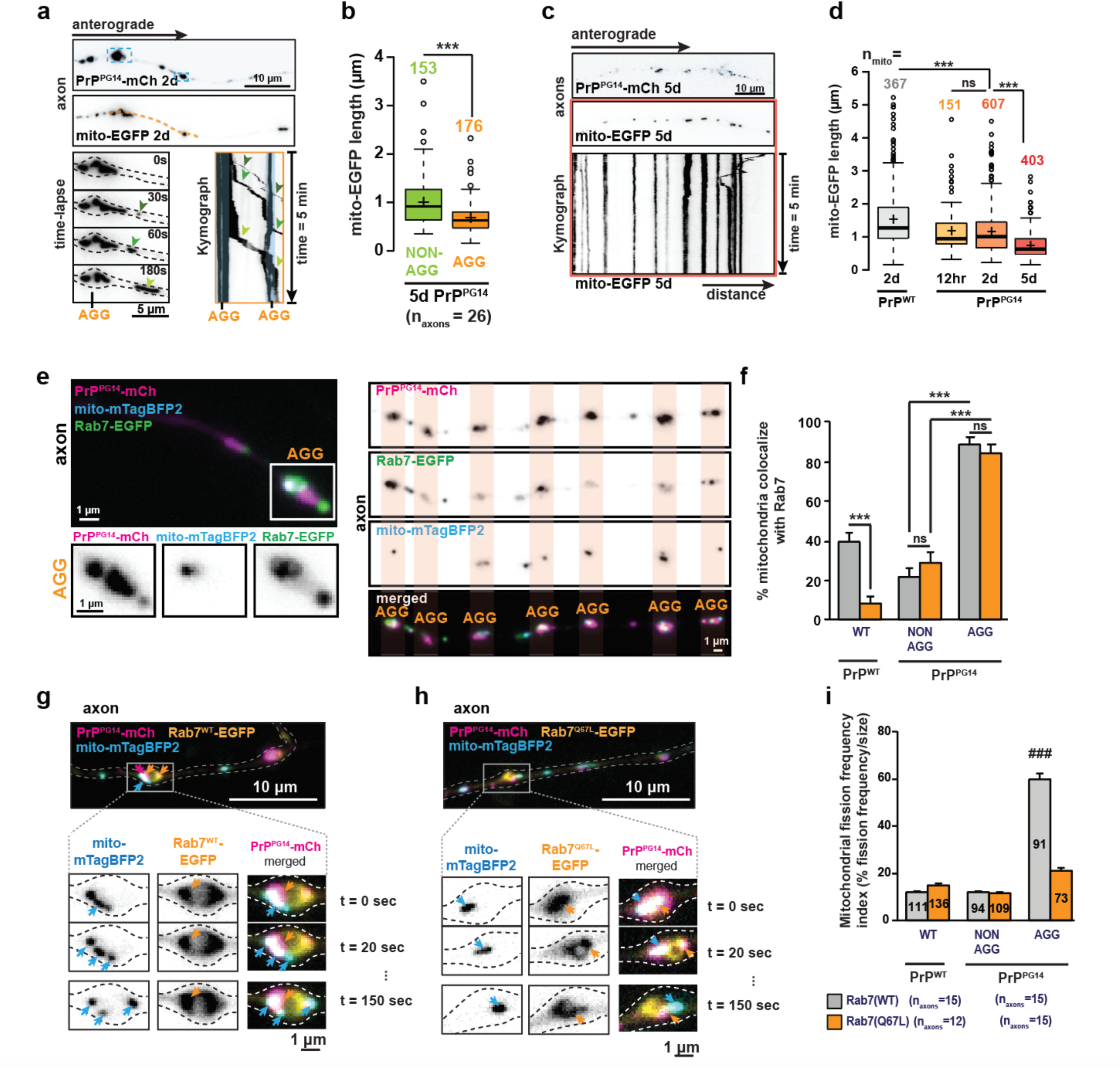
Rab7-mediated mitochondrial fission at PrP^PG14^ aggregate sites. **a** First-frame images of axons co-expressing PrP^PG14^-mCh and mito-EGFP for 2 d. The yellow dotted line indicates an imaging region of time-lapse (bottom left) and kymograph (right). **b** Quantitation of mito-EGFP length at AGG vs NON-AGG sites at 5 days post-PrP^PG14^-mCh expression. **c** First-frame images of axons co-transfected with mito-EGFP and PrP^PG14^-mCh for 5 days (top two panels) and corresponding kymographs (bottom panel) generated from time-lapse movies (5 minutes, 300 frames, 1 frame/sec). **d** Quantitation of mito-EGFP length co-transfected with PrP^WT^-mCh for 2 days or PrP^PG14^-mCh for 12 hours, 2 days, and 5 days. e Axons co-expressing PrP^PG14^-mCh, Rab7-EGFP, and mito-mTagBFP2. Insets and highlights indicate AGG sites. **f** Quantitation of percent mito-mTagBFP2 colocalize with Rab7(WT/Q67L)-EGFP at WT, NON-AGG, or AGG sites. **g** First-frame images of axons co-expressing PrP^PG14^-mCh, Rab7^WT^-EGFP, and mito-mTagBFP2 and representative frames from time-lapse videos at WT or AGG sites (insets). **h** First-frame images of axons co-expressing PrP^PG14^-mCh, Rab7^Q67L^-EGFP, and mito-mTagBFP2 and representative frames from time-lapse videos at WT or AGG sites (insets). **i** Quantitation of mitochondrial fission frequency index (% of fission frequency / mitochondrial length). Box plots were shown with Tukey definition; (+) indicates the mean. All bar graphs are shown as mean ± SEM. ***p<0.001, **p<0.01, *p<0.05, ns = not significant; **b, d** Kruskal-Wallis test. **f, i** Tukey’s multiple comparison test. See also figure S4.

Previous studies using ultrathin serial-sectioning SEM (S3EM) to visualize the ultrastructure of PrP^PG14^ axonal swellings showed mitochondria and endolysosomes trapped within PrP^PG14^ endoggresome AGG sites and making close contacts^17^. Endolysosomes marked with Rab7, a small GTPase that binds to endosomal membranes upon GTP hydrolysis^55^, have been shown to regulate mitochondrial fission via direct contact with mitochondria, to promote fission at contact sites^56^. To investigate the mechanistic basis of enhanced mitochondrial fission within AGG sites, we first used live high-resolution HILO imaging to test whether Rab7 associated with mitochondria within PrP^PG14^ endoggresome AGG sites. Neurons were co-transfected with mito-mTagBFP2 plus PrP^WT^-mCh or PrP^PG14^-mCh plus Rab7(WT)-EGFP or Rab7(Q67L)-EGFP. The latter is a constitutively active mutant that cannot undergo GTP hydrolysis and thus cannot be released from endosomal membranes^57^, and that has been shown previously to inhibit mitochondrial fission^58^. Colocalization analysis showed that mito-mTagBFP2 highly colocalized with Rab7(WT/Q67L)-EGFP at AGG sites vs NON-AGG sites of PrP^PG14^-mCh endoggresome-rich axons versus in control PrP^WT^-mCh endoggresome-free axons at 2 days post-transfection (**Fig. 6e,f, Supplementary Fig. 4a**). HILO high-resolution imaging revealed mitochondrial (mito-mTagBFP2) fission occurred within PrP^PG14^-mCh AGG swelling sites in neurons expressing Rab7(WT)-EGFP (**Fig. 6g**). Importantly, fission did not occur as frequently in PrP^PG14^-mCh AGG swelling sites of neurons also expressing the constitutively active Rab7(Q67L)-EGFP (**Fig. 6h**). To quantitate these observations, we calculated a mitochondrial fission index as the frequency of fission normalized to mitochondrial lengths. These data indicate that the frequency of mitochondria fission at AGG sites was significantly higher than either fission in PrP^WT^ axons or in NON-AGG regions of axons of neurons expressing PrP^PG14^-mCh (**Fig. 6i**). Mitochondria also underwent fission in axons of neurons co-expressing Rab7(WT)-EGFP but not in those overexpressing Rab7(Q67L)-EGFP (**Supplementary Fig. 4a, b**), consistent with an established role of Rab7 GTPase hydrolysis in mitochondrial fission in healthy cells^56^. Altogether, our observations indicate that PrP^PG14^ AGG swellings act as hubs for organelle-organelle contact hotspots that mediate mitochondrial remodeling via promoting its fission through Rab7 GTPase activity.

### Dysfunctional mitochondria are retained at PrP^PG14^ aggregate sites

Mitochondrial dysfunction occurs at early and late stages of neurodegeneration, including in prion diseases^59–61^. Mitochondrial membrane potential (ý4′m) is central to neuronal health, and both increased and decreased membrane potential have been associated with impairments to neuronal mitochondrial quality control in familial and infectious prion disorders^61, 62^. We hypothesized that the fragmented mitochondria that accumulate at PrP^PG14^ AGG sites become dysfunctional resulting in a defective ability to retain their membrane potential. To test this, we measured the membrane potential of mitochondria within individual PrP^PG14^ endoggresome AGG swellings and compared it to that at NON-AGG sites or to mitochondrial membrane potential in axons of neurons expressing PrP^WT^-mCh controls. Neurons co-transfected with PrP^PG14^-mCh or PrP^WT^-mCh and Rab7(WT)-EGFP -which we showed potentiates mitochondrial fission at AGG sites (**Fig. 6g**)-, were loaded with TMRE (tetramethylrhodamine, ethyl ester), a cell permeant dye that accumulates and labels active mitochondria, and that is not sequestered within depolarized or inactive mitochondria^63, 64^. Quantitation of TMRE intensity at mito-EGFP showed a significant reduced TMRE signal that mito-EGFP at the AGG sites had (**Fig. 7a, b**), and the percentage of mito-EGFP that could retain TMRE dropped significantly, compared to mito-EGFP in NON-AGG and WT control sites (**Fig. 7c**). Thus, PrP^PG14^ endoggresome-mediated mitochondrial transport dysfunction and fragmentation are associated with compromised mitochondrial integrity within PrP^PG14^ axonal swellings.

**Fig. 7:**
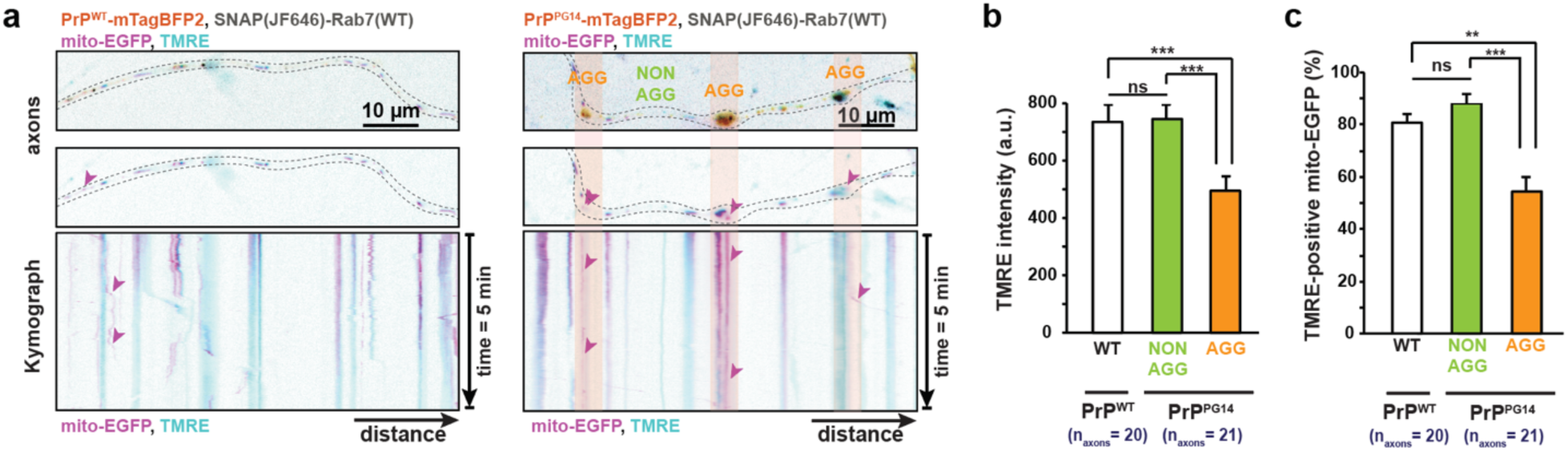
Dysfunctional mitochondria accumulates within PrP^PG14^ aggregate sites. **a** Representative inverted color first frames of axons (top two panels) and kymographs (bottom panel) generated from time-lapse movies (5 minutes, 300 frames, 1 frame/sec) of PrP^WT^- or PrP^PG14^-mTagBFP2, co-transfected with SNAP(JF646)-Rab7 and mito-EGFP and treated with 25 nM of TMRE. Arrowheads point to TMRE-negative mito-EGFP. **b** TMRE intensities per mito-EGFP in WT, NON-AGG, or AGG regions. **c** Percent mito-EGFP that are TMRE-positive. All bar graphs are shown as mean ± SEM. ***p<0.001, **p<0.01, *p<0.05, ns = not significant; **b, c** Tukey’s multiple comparison test.

## DISCUSSION

Our previous work identified an endolysosomal pathway in mammalian axons call **<u>a</u>**xonal **<u>r</u>**apid **<u>e</u>**ndosomal **<u>s</u>**orting and **<u>t</u>**ransport-dependent **<u>a</u>**ggregation (ARESTA), as a principal driver of the initial stages of formation of enlarged misfolded mutant PrP aggregates inside endolysosomes (called endoggresomes) in axons, including of PrP^PG14^ ^17^. ARESTA drives the formation of PrP^PG14^ endoggresomes via the homotypic sorting and fusion of endolysosomes containing PrP^PG14^, as they traffic along axons^17^. Yet, how enlarged mutant PrP endoggresomes that distend the axon lead to axonal pathologies (axonopathies) and eventually to axonal degeneration, has remained unknown. In this study, we modeled axonal PrP^PG14^ aggregation in a cultured primary murine hippocampal neuronal system over time, to unveil the mechanisms of localized cytoskeletal disruptions and organelle accumulations within local axonal swellings where PrP^PG14^ endoggresomes form. Using high-and super-resolution light microscopy, including single-particle micro-movement analyses of cargoes, and correlative light and EM imaging, we dissected the mechanisms of local impairments that develop at the subcellular and ultrastructural levels within aggregate swellings in axons. This detailed characterization revealed that PrP^PG14^ endoggresome swelling sites represent micro-hubs of toxicity along axons, where cascades of defects ensue at very early stages after expression of PrP^PG14^. These localized micro-phenotypes include cytoskeletal remodeling, anterograde ‘transport jams’ due to altered architecture of MT tracks that kinesin-1 and kinesin-3 cargoes mobilize on, and the sequestration of mitochondria, endolysosomes, and molecular motors at swelling sites (**Fig. 8**). We propose that following this cascade of events, PrP^PG14^ axonal swelling sites become ‘organelle-organelle contact hotspots’ that drive the close contact between mitochondria and active Rab7-positive endolysosomes, to promote mitochondrial remodeling (via enhanced fission), and thus compromise mitochondrial functional integrity in axons (**Fig.8**).

**Fig. 8:**
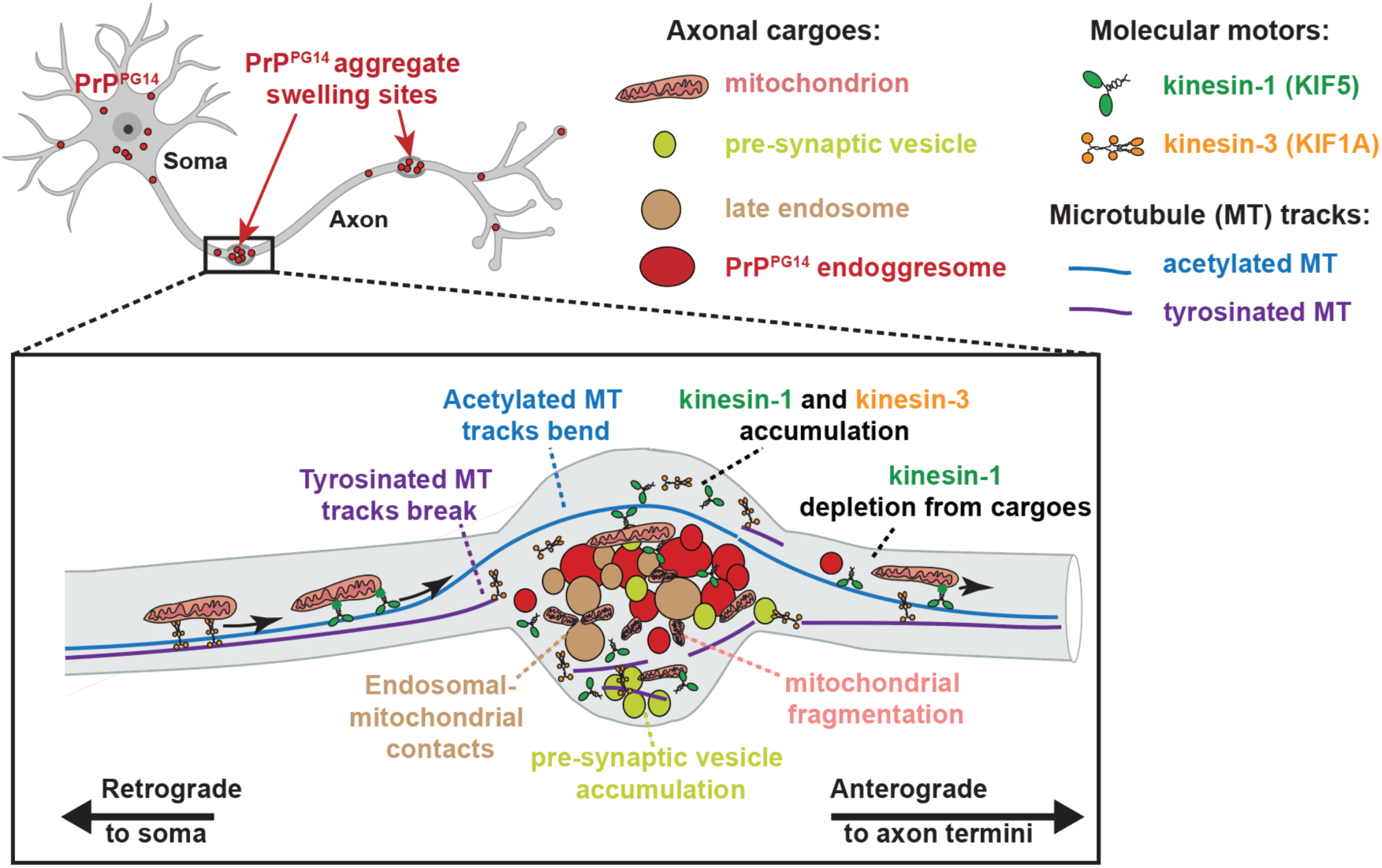
Working model of the mechanism of cytoskeletal-organelle remodeling at PrP^PG14^ endoggresome swelling sites. PrP^PG14^ endoggresomes form at various sites along mammalian axons where they distend axons into swellings. At endoggresome sites (1) acetylated MT tracks bend and tyrosinated MT tracks break at these swelling sites, (2) kinesin-1 and -3 are sequestered and are depleted from mitochondria that mobilize through these sites, and (3) axonal transport of mitochondria, pre-synaptic and PrP^PG14^ vesicles is selectively impaired in the anterograde direction, and (4) PrP^PG14^ endoggresomes trap Rab7-endolysosomes and mitochondria, enhancing the close contact between these organelles, leading to elevated mitochondrial fission and dysfunction.

Previous studies showed that MT stabilization is conferred by regulation of acetylation vs tyrosination, two PTMs that elicit MTs to bend or break under mechanical stress^43^. Our data are consistent with these observations, as acetylated MTs curve while tyrosinated MTs break at PrP^PG14^ endoggresome swelling sites (**Fig. 3**), suggesting that enlarged PrP^PG14^ endoggresomes exert mechanical stress from within axons. Our data further suggest that breakage of tyrosinated MTs at swelling sites could be an important initiating event that precedes the cascade of downstream deficits, including transport defects and motor and organelle accumulations that are observed within localized swelling regions. Tyrosinated MTs are preferred by kinesin-3 (KIF1), and this motor is used by mitochondria, pre-synaptic vesicles, and PrP vesicles (R. Chassefeyre, pers. comm.) to move toward the synapse^65–67^. On the other hand, kinesin-1 (KIF5) prefers bundled and stable acetylated MTs^46, 68^ and is also a motor for the movement of mitochondria, pre-synaptic vesicles, as well as of wild-type and mutant PrP vesicles^17, 34^ along axons^65–67^. Thus, kinesin-3 transport is expected to be directly affected by broken tyrosinated MTs, while bending acetylated MTs likely sustain the transport of kinesin-1 cargoes, including of mitochondria, through axonal swollen regions. As it is unclear how kinesin-1 and kinesin-3 coordinate activities when bound simultaneously to a cargo^47, 69^, it is possible that complex interactions between these motors and MT tracks dictate the final outcome of motility of mitochondria within and through these sites.

The global deficits observed primarily to the anterograde axonal transport of three different cargo—mitochondria, pre-synaptic and mutant PrP vesicles—point to disruptions to generalized motility pathways in axons of neurons expressing PrP^PG14^, that are likely dictated at least in part, by the observed alterations to subsets of MTs with different PTMs. However, MT defects might not be responsible for the very early transport defects first observed for the anterograde movement of vesicles carrying PrP^PG14^ at 12 hours post-transfection, when only very few endoggresomes have formed (**Supplementary Fig. 1**), and when presumably the MT cytoskeleton is intact. These observations suggest that other even earlier deficits ensue in axons. A possibility is that the previously observed initial active and dynamic sorting of endolysosomes containing PrP^PG14^ into fused enlarged organelles^17^ could inhibit transport of those endolysosomal vesicles during the process of forming larger endoggresomes. As mitochondrial movement is unaffected at the 12 hour earlier time point, our data suggest that mitochondrial motility impairments represent local traffic disruptions that result from micro-transport deficits that start at AGG sites following the formation of PrP^PG14^ endoggresomes, thus implicating PrP^PG14^ endoggresomes directly in the initiation of these local transport deficits.

The observation that anterograde, but not retrograde transport of PrP^PG14^ vesicles, pre-synaptic vesicles, and mitochondria, was impaired in axons of neurons expressing PrP^PG14^, was unexpected. Previous work showed that disruptions to kinesin-mediated anterograde movement of various cargoes in axons also impaired dynein-mediated retrograde transport because these opposing motors often associate simultaneously to the same cargo to coordinate activities^34, 70^. Both kinesin-1 and dynein can associate to mitochondria simultaneously^71^, thus a possibility is that mitochondria moving in the retrograde direction by dynein, the principal retrograde motor in axons^72^, utilize intact acetylated MTs tracks. Previous *in vitro* studies showed that dynein associates to tyrosinated MTs tracks to initiate is movement, but that subsequently dynein can continue translocating on detyrosinated MTs^73^. Thus, it could be that dynein initiates movement on broken tyrosinated MTs in axons expressing PrP^PG14^, but continues movement on acetylated (detyrosinated) MTs, and this possibility merits further exploration. Differences in anterograde versus retrograde transport impairments could also stem from differences in the intrinsic properties of kinesin and dynein proteins. Several studies using *in vitro* motility and biophysical assays showed that a single dynein molecule could take side-steps around MT tracks to avoid obstacles in dense cytoplasmic environments during cargo transport^74, 75^. Kinesin-1, on the other hand, walks linearly on single protofilament tracks and works in teams to avoid obstacles^75^.

Thus, in the normal crowded environment of axons, which are accentuated by the presence of enlarged PrP^PG14^ endoggresomes, dynein-driven cargoes might be capable of navigating through dense cytoskeletal networks while kinesin-1 molecules might not. Notably, a previous study in an PrP^Sc^-infected mouse model suggested that the MT network and dynein-mediated retrograde transport were not directly affected, but instead, they proposed that motor-cargo associations might be the target of infectious prions prior to clinical onset *in vivo*^76^. While it is unclear whether mechanisms of axonal impairment in infectious PrP^Sc^ models are similar to those in familial forms of prion disease, our data is consistent with this scenario, as we showed that motor-cargo associations were impaired and that kinesin-1 detachment and accumulation at PrP^PG14^ aggregate sites depend on the presence of the cargo binding domain of kinesin-1 (**Fig. 4**).

In prion diseases, ultrastructural studies of human patient brains with prion disease have revealed the accumulation of misfolded PrP within dystrophic neurites inside endolysosomes^18– 21, 77, 78^. Other complex structures have been observed to co-exist with aggregates within swollen neurites, and are also considered common features of prion pathology, including autophagic vacuole-like membrane-bound organelles, lysosomal electron-dense bodies, enlarged endolysosomes, and membraneless electron-lucent areas of cytoplasm^18, 20, 77^. Some of these neuronal phenotypes have also been observed in cell lines infected with PrP^Sc^, as well as in brains of rodent models of familial and sporadic human PrP^Sc^ disease ^18, 77–81^. The presence of dystrophic swellings in axons is also a common and early feature in brains of patients in various neurodegenerative pathologies including prion diseases, AD, PD, and HD^3, 14, 82, 83^. In AD, dystrophic axons are also comprised of enlarged endosomes, and these lesions are considered amongst the earliest neuronal lesions identified thus far^84, 85^. These observations highlight the importance of understanding the mechanisms of axonotoxicity as imparted by the presence of intra-axonal misfolded protein aggregates in these diseases.

In the familial PrDs, the formation of axonal misfolded PrP aggregates contributes to neurodegeneration, as reduction of axonal aggregates by modulating its biogenesis pathway rescues neuronal function^17^. To what extent modulation of endolysosomal pathways contributes to the formation of aggregates in other neurodegenerative disorders is unknown. However, as misfolded proteins like tau, α-synuclein, APP and Huntingtin are associated at least in part with endolysosomes and undergo sorting and trafficking^86–88^, it will be key to determine whether the defects to the cytoskeleton and micro-domain impairments observed in PrP^PG14^-expressing neurons also participate in axonotoxicity in the rest of the proteinopathies. The subcellular lesions identified in this study provide new pathways that can be therapeutically targeted for their amelioration and for the rescue of axonal lesions that are the primary vulnerability targets in the proteinopathies.

## METHODS

### DNA constructs

PrP^WT^-EGFP and PrP^PG14^-EGFP in the MoPrP.Xho vector^89^ were a gift from David A. Harris^16^. PrP^WT^-mCh, PrP^PG14^-mCh, PrP^WT^ -mTagBFP2, PrP^PG14^-mTagBFP2 fusions were cloned by assembly PCR to replace EGFP as described previously^17^. Synaptophysin-YFP (SYP-YFP) in cEGFP vector was a gift from Ann Marie Craig^90^. Mito-EGFP in pcDNA3 vector was a gift from Christine Vande Velde^91^. Mito-mTagBFP2 was cloned by replacing EGFP in mito-EGFP with mTagBFP2 sequence in pcDNA3.1 from mTagBFP2-Lysosome-20 (Addgene #55308). KIF5C(1-560)-2xmCh-EF(C) (Addgene #61664) and KIF5C(1-560)-mCit (Addgene #61676) were a gift from Kristen Verhey^92^. 2xmCh-KIF5C in pcDNA3.1 was cloned by inserting in-frame KIF5C full length from EGFP-KIF5C (a gift from Anthony Brown) 2xmCh sequence from KIF5C(1-560)-2xmCh-EF(C) into pcDNA3.1 using InFusion Cloning kit (Takara). EGFP-Rab7A (Addgene #28047) and EGFP-Rab7A Q67L (Addgene #28049) were a gift from Qing Zhong. SNAP-Rab7A and SNAP-Rab7A Q67L were made by cloning Rab7A sequence from EGFP-Rab7A and EGFP-Rab7A Q67L plasmids into pSNAP-tag (m) Vector (Addgene #101135). Soluble BFP (mTagBFP2) in pcDNA3.1 was derived from mTagBFP2-Lysosome-20 (Addgene #55308). Soluble EGFP (pEGFP-C1 plasmid) was a gift from Hilal Lashuel. Soluble mCherry (pmCherry-N3 plasmid) was cloned from pmCherry-mito-7 (Addgene #55102).

*Kif1a* and *scrambled* shRNAs were part of a kinesin lentiviral mini-library, in a pLL3.7 GW lentiviral vector with Gateway entry modifications and a mCherry co-expression marker (unpublished data). The knockdown efficacy of *Kif1a* shRNA was evaluated by Western blot and qPCR from lysates of cultured hippocampal neurons transduced with lentivirus containing *Kif1a* shRNA-expressing plasmids (unpublished data).

### Antibodies

The following primary antibodies were used for immunofluorescence (IF): mouse anti-KHC (clone H2; MAB1614; recognizes primarily KIF5C^34^) (1:100, EMD Millipore); goat anti-KLC1 (V-17) (1:100, Santa Cruz Biotechnology); mouse anti-KIF1A (16/KIF1A) (1:100; BD Transduction Laboratory); mouse anti-alpha-tubulin (DM1A) (1:2000; Sigma-Aldrich); mouse anti-Acetylated tubulin (1:400; Sigma-Aldrich); rat anti-Tyrosinated tubulin [YL1/2] (1:1000; Abcam). The concentration of Goat anti-KLC1 (V-17) was validated previously for sub-pixel colocalization analysis^34^. The concentration of mouse anti-KHC and mouse anti-KIF1A were also validated for sub-pixel colocalization analysis (data not shown). All tubulin antibodies were validated previously to be suitable for IF for super-resolution imaging^48^.

The following secondary antibodies were used for IF: Donkey anti-goat IgG (H+L) Alexa Fluor 488 (1:200; ThermoFisher; A-11055); Goat anti-mouse Atto647N (1:200 Sigma); Goat anti-rat Atto594 (1:200; Rockland Immuno); Alexa Fluor 647 Phalloidin in methanol (1:40; Invitrogen)

### Cultured primary mouse hippocampal neurons

Primary mouse hippocampal neurons were prepared as described previously^17, 34, 93^.

Briefly, hippocampi were dissected from 0 to 2-day-old mouse neonates, treated with Papain (Worthington) for 15 minutes, and dissociated by seven to ten cycles of aspiration through micropipette tips. Dissociated neurons were resuspended in Neurobasal-A medium containing 2% B-27 (Gibco) and 0.25% GlutaMAX supplements (Gibco). Neurons were plated in 24-well plates containing #1.5 1-mm round coverslips (Neurovitro) and were maintained at 37°C and in a 5% CO_2_ atmosphere up to 12 DIV for imaging.

### Transfection and live imaging of hippocampal neurons

Neurons were transfected at between 4-7 DIV. We used 2 µL of Lipofectamine 2000 (Invitrogen) and up to 1.6 µg total of DNA per well in 24 well plates. Transfection rates of cultured hippocampal neurons were ∼5%, which enabled imaging of individual axons. Neurons were imaged using a Nikon Ti-E Perfect Focus inverted microscope equipped with a total internal reflection fluorescence (TIRF) setup, with an Andor iXon + DU897 EM Camera, and a 100X/1.49 NA oil objective. A 405-nm, 488-nm, 561-nm, and 640-nm laser lines were used to detect BFP, GFP, mCh, and JF646 fluorescence respectively, at an angle for optimal pseudo-TIRF/HILO acquisition. Movies of vesicular transport were 15 seconds long and collected at a frame rate of 10 frames/sec (10 Hz). Movies of mitochondrial transport were 5 minutes long and collected at a frame rate of 1 frame/sec (1 Hz). Cultured neurons were maintained at 37°C and 5% CO_2_ throughout the total imaging period.

### Quantitative analysis of cargo motility in whole axons

Analysis of cargo motility was performed by using *KymoAnalyzer*, an ImageJ macros package specifically designed to track the movement of particles in axons^35^. Briefly, kymographs (distance-time projections) were generated from time-lapse movies. Particle trajectories were manually assigned from the kymograph images. Track and segment-related parameters were automatically calculated. Detailed definitions of all the transport parameters analyzed in this study were described previously^35^.

### Single-particle tracking (SPT) of mitochondria

SPT of mito-EGFP was done using *TrackMate*, a Fiji (ImageJ) plugin^94^. In brief, Laplacian of Gaussian (LoG) filter, median filter, and sub-pixel localization were applied to detect the blobs with diameter between 1-3 µm. Auto-thresholding was applied on the detected feature quality to restrict number of spots. The hyperstack displays contain tracks analyzed by LAP Tracker algorithm^95^ with color codes based on parameters such as median velocity or displacement. Frame-to-frame linking maximum distance was set to 15 µm. Gap closing, segment splitting and merging were disabled to take into account all movement instances as independent.

### Quantitative analysis of local cargo motility disruption

Local transport disruption analysis was performed manually. AGG sites are defined as a 10-µm window along axonal length containing particles that had fluorescence intensities above the average grey value of five traveling fluorescently labelled PrP^PG14^ cargos. NON-AGG sites are defined by a 10-µm window along axonal length that do not contain particles that had fluorescence intensities above the average grey value of five traveling fluorescently labelled PrP^PG14^ cargos. Particle trajectories were categorized into 3 groups. Pass tracks on kymographs are defined as diagonal tracks that cross AGG/NON-AGG sites without any change in slope. Pause/Slowdown tracks are defined as diagonal tracks that cross AGG/NON-AGG sites with any change in slope. Stall tracks are defined as stationary tracks with slope equal to 0 at AGG/NON-AGG sites.

### Immunofluorescence and imaging of fixed neurons

Neurons were fixed with 4% PFA containing 4% sucrose during 30 minutes at 37°C. Neurons were washed 3 times in PBS before permeabilization in 0.1% TritonX-100 in PBS. After another 3 washes in PBS, neurons were incubated in a blocking solution containing 3% IgG-free bovine serum albumin, 5% Normal Goat/Donkey serum (Jackson ImmunoResearch) in PBS during 30 minutes at room temperature. Primary antibodies were incubated in blocking solution overnight at 4°C. After 3 washes in PBS, secondary antibodies were incubated in blocking solution during 1 hour at room temperature. Neurons on the coverslips were mounted onto glass slides using ProLong Diamond antifade reagent (Thermo Fischer) and imaged within 2-3 weeks post-fixation.

### Quantitation of kinesin-1 and kinesin-3 fluorescence intensity

KHC was detected with a validated antibody that recognizes primarily KIF5C^34^. The analysis was done in Fiji (ImageJ). Briefly, normalized fluorescence intensity from AGG, NON-AGG, or WT sites was calculated from averaged gray values of antibody staining in those regions subtracted background fluorescence per image. Regions were selected by unbiased method, by copying ROIs from PrP^PG14^/PrP^WT^ channels to antibody channels, and the gray values were obtained in a batch analysis. The fluorescence intensity from the whole dataset then was plotted in the OriginLab software (https://www.originlab.com/).

### Sub-pixel colocalization analysis

Single-particle detection with sub-pixel precision was performed as described previously^34, 53^. In brief, all primary and secondary antibodies were validated for specificity and determined saturation curve. Tetraspeck beads were used to perform registration and to correct chromatic aberration between channels. Secondaries-only controls were included in every experiment. Raw fluorescence images were processed in MATLAB for single-particle detection. To determine motor colocalization on mitochondria in Fiji (ImageJ), ROIs were drawn around mito-EGFP signal and transferred to particle-detected images output from MATLAB, and number of motor clusters were counted manually.

### FRAP

FRAP of KIF5C(1-560)-mCit was performed at the axon terminals. Briefly, A 405-nm laser was collimated and positioned in the middle of the field of view. Laser power and exposure time were determined empirically to ensure complete bleaching and avoid phototoxicity. For the acquisition, 5 images were collected before photobleaching (at 1 frame/sec; 1 Hz), immediately followed by a bleaching of 1-3 seconds, and 300 images were collected after photobleaching (at 1 frame/sec; 1 Hz). Before and after images were combined automatically in Nikon Elements software, and fluorescence intensity plots over time were analyzed in Fiji (ImageJ). Half-life (t_1/2_) was calculated by fitting one-component exponential curve, using FRAP profiler ImageJ plugin (http://worms.zoology.wisc.edu/ImageJ/FRAP_Profiler_v2.java).

### SRRF

SRRF processing was done using NanoJ-SRRF ImageJ plugin (https://github.com/HenriquesLab/NanoJ-SRRF). Briefly, 100 frames were collected at 50 frames/sec; 50 Hz. SRRF parameters were set as default (ring radius = 0.5; radiality magnification = 5; axes in ring = 6). Drift correction was applied.

### STORM sample preparation and imaging

Hippocampal neurons were plated on MatTek glass bottom dishes (acid pre-washed according to N-STORM preparation protocol) and were transfected with PrP^PG14^-mCh at DIV7. By 2 days post-transfection (DIV9), neurons were fixed in 3% PFA, 0.1% Glutaraldehyde (GA), 10% sucrose in PBS. Neurons were permeabilized in 0.1% Tx-100 for 8 minutes. Then, neurons were incubated in 0.5 µM of Phalloidin-AF647 overnight at 4°C. Neurons were washed and kept in PBS at 4°C to be imaged within 2 days post-staining.

STORM buffer was prepared according to STORM Protocol-Sample Preparation (Nikon, 2013), which included 50 mM TRIS, 10 mM NaCl pH 8, 0.5 mg/mL glucose oxidase, 40 µg/mL catalase, 10% glucose, and 100 mM β-mercaptoethanol (BME). STORM buffer was applied on the MatTek dish sample and sealed tightly before mounting on the microscope. More than 10,000 raw images were captured with Nikon N-STORM acquisition settings using 640-nm laser and 100X/1.49NA Objectives. STORM images were reconstructed using ThunderSTORM ImageJ plugin^96^. The display mode was based on an average shifted histogram approach, which provides similar results as conventional Gaussian rendering. All dSTORM images collected had sufficient blinking of AlexaFluor647 over 10,000 frames during acquisition.

### Membrane-extraction SEM (ME-SEM) preparation and imaging

Neurons were plated on sterile MatTek dish containing gridded coverslip (https://www.mattek.com/store/p50g-2-14-fgrd-case/). Sterile custom-made PDMS molds were temporary attached on the dish to create smaller chamber with the same dimension (volume & surface area) as 24-well plate. Neurons were plated at the concentration of 2 hippocampi into 12 chambers (1:12) or lower and transfected with PrP^PG14^-mCh at DIV7 following previously described protocol. Membrane extraction was performed 2 days post-transfection (DIV9). Live neurons were imaged with 20X/0.45NA objectives with phase contrast mode and 561-nm laser to find transfected neurons and its grid coordinates. Then, live neurons were treated with a membrane extraction solution containing 1% Triton X-100, 100 mM PIPES (pH 7.2), 4% sucrose, 1 mM MgCl_2_, 10 μM taxol (Invitrogen) and 10 μM phalloidin (Sigma)^97^. After 20 minutes of gentle rocking, the neurons were washed three times with PBS with gentle pipetting during each wash to remove membrane debris from the sample. Neurons were then fixed with 2% glutaraldehyde, 4% paraformaldehyde [Electron Microscopy Sciences (EMS), Hatfield, PA, USA], 140 mM NaCl, 2 mM CaCl_2_ and 1 mM MgCl_2_ in 0.1 M HEPES (pH 7.2) for 30 minutes. Samples were transferred and post-fixed with 1% OsO4 (EMS) and 1% tannic acid (EMS) for 1 hour each, dehydrated with ethanol and critical point-dried from CO_2_ (CPD 020 Balzers-Tec), and the coverslips were coated with a thin carbon layer (8 nm) (BAF 300, Balzers)^97^ for imaging on Zeiss SIGMA Variable Pressure Field Emission Gun Scanning Electron Microscope.

Fluorescence images and SEM images were aligned manually on Adobe Illustrator.

### Figure plotting and statistical analyses

Histograms were plotted in MATLAB (MathWorks), and Gaussian mode mixtures were generated using the Interactive Peak Fitter (ipf.m). Optimal fits were selected empirically based on optimal R^2^ values. Box plots were plotted using the web-tool BoxPlotR (http://shiny.chemgrid.org/boxplotr/). Cumulative frequency distribution graphs were plotted using GraphPad Prism. Statistical analyses were performed using Microsoft Excel, GraphPad Prism, and MATLAB. Briefly, we first checked if the parameters were following a normal distribution using the Lilliefors test. For the parameters following a normal distribution (distribution of cargo populations, densities, etc.), we performed a Student’s t-test or Tukey’s multiple comparisons. Most of the other parameters that did not follow a normal distribution (velocities, run lengths, fluorescence intensity, etc.) were subjected to appropriate non-parametric tests. Differences between two medians were compared using the Wilcoxon rank-sum test, while pair-wise differences between three groups were compared with Kruskal-Wallis test. Differences between two population distribution were compared using Kolmogorov-Smirnov (K-S) test.

### TMRE assay

Neurons for the TMRE assay were grown on the coverslip and transfected with 400 ng of SNAP-Rab7, 800 ng of PrP^WT^-mTagBFP2 or PrP^PG14^-mTagBFP2, and 200 ng of mito-EGFP. At 2 days post-transfection, neurons were treated with 25 nM TMRE (Invitrogen) in growth media for 20 minutes at 37°C as described previously^64^. Neurons were then washed 1 time in pre-warm PBS. Then, neurons were treated with 100 nM JF646-SNAP-tag ligands in growth media for 30 minutes at 37°C as described previously^98^. Neurons were washed 1 more time in warm PBS and put back in the growth media to perform live imaging immediately. 568-nm laser was used to detect TMRE signal, and 3-color simultaneous time-lapse imaging were collected at 0.5 frame/sec (0.5 Hz) on the OptoSplit III (Andor).

## ACKNOWLEDGEMENTS

We thank Isabelle C. Romine for preliminary data collection; George Campbell for antibody validation and sub-pixel motor colocalization optimization; Tong Zhang for Airyscan microscopy training and data collection; Kathryn Spencer for STORM imaging training; Michael Petrascheck, Jeff Kelly, Kayal Madhivanan, David Soriano Castell, Andre Leitao, and current and past members of the Encalada lab for advice and constructive feedback. This work was supported, in part by NIH/NIA R01AG049483 and NIH/NIA R01AG076745-01 grants (to S.E.E.), the Glenn Foundation for Medical Research Award for Research in Biological Mechanisms of Aging (to S.E.E.), New Scholar in Aging Award from the Lawrence Ellison Foundation, Baxter Family Foundation award (to S.E.E.), T.C. was supported by a Royal Thai Government Scholarship from the Development and Promotion of Science and Technology Talents Project (DPST) and the Skaggs Graduate School of Chemical and Biological Sciences and Scripps Research, George E. Hewitt Foundation for Medical Research Postdoctoral Fellowship (to R.C.), Dorris Neuroscience Center Scholar Award (A.V.), Larry L. Hillblom Foundation Inc. (to U.M. and S.W.N.), Chan-Zuckerberg Initiative (to U.M.), Waitt Foundation Core Grant application NCI CCSG (CA014195) (to U.M., S.W.N., and L.R.A.), NSF NeuroNex Award no. 2014862 (to U.M. and S.W.N. and NIH/NIDCD R21 award DC018237 (to U.M. and L.R.A.).

## Author Contributions

Conceptualization: T.C. and S.E.E. Methodology: T.C., A.V., R.C., N.S., S.W.N., L.R.A., U.M., and S.E.E. Investigation: T.C., A.V., R.C., N.S., S.W.N., L.R.A.Validation: T.C., A.V., S.W.N., and L.R.A. Supervision: S.E.E. and U.M. Formal analysis: T.C., A.V., R.C., N.S., S.W.N., U.M., and S.E.E. Writing (original draft): T.C and S.E.E. Writing (review and editing): S.E.E., T.C.

## MATERIALS AND CORRESPONDANCE

Should be addressed to S.E.E. at

## SUPPLEMENTAL INFORMATION

**Supplementary Video 1.**
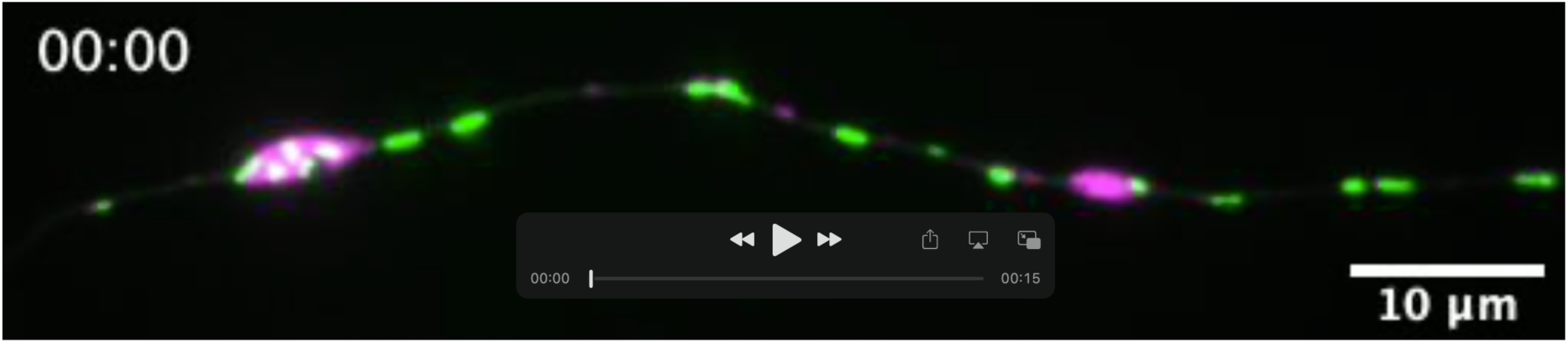
Mitochondrial transport disruption and mitochondrial fission at PrP^PG14^ aggregate sites. A 5-minute movie of an axon co-expressing PrP^PG14^-mCh (magenta) and mito-EGFP (green) showing transport disruption (a swelling site on the right) and mitochondrial fragmentation (a swelling site on the left). Movement towards the right side of the video is anterograde direction. (Related to **Fig. 1 and Fig. 6**)

**Supplementary Fig. S1:**
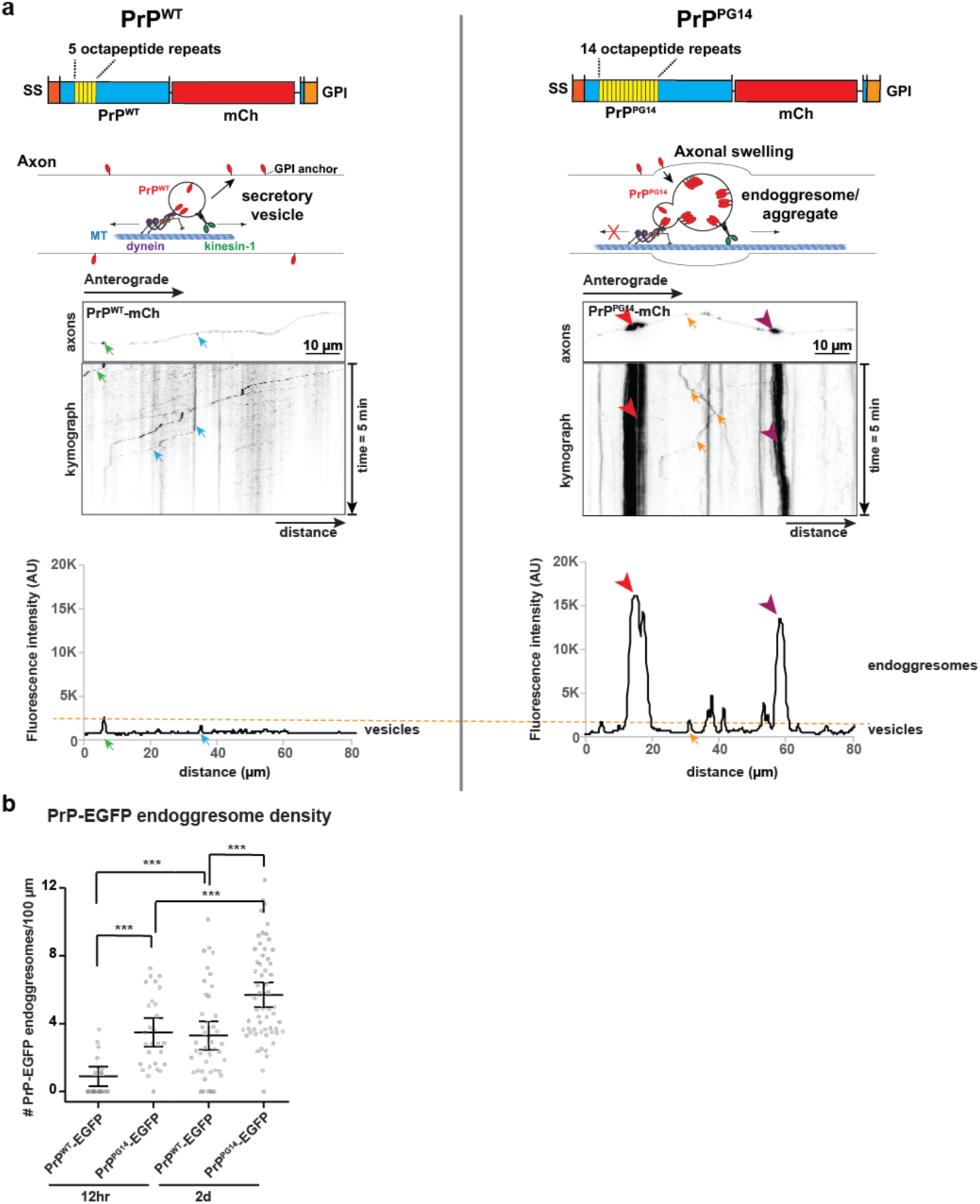
PrP^PG14^ forms aggregates inside axons. **a** Schematic of PrP^WT^-mCh and PrP^PG14^-mCh constructs (top panels; SS: signal sequence, GPI: GPI anchor), and representative images of axons first frame of a time-lapse movie showing PrP^WT^-mCh and PrP^PG14^-mCh vesicle transport in hippocampal axons of cultured neurons. Corresponding kymographs are shown below first movie frames. Fluorescence intensity profiles obtained from line scan plots (bottom panels), highlight key differences between PrP^PG14^-mCh and PrP^WT^-mCh control. Arrows point to vesicles. Arrowheads point to aggregates. **b** Quantitation of density of PrP-EGFP endoggresomes. The graph is shown as mean ± SEM with all datapoints. ***p<0.001, **p<0.01, *p<0.05, ns = not significant. Tukey’s multiple comparison test.

**Supplementary Fig. S2:**
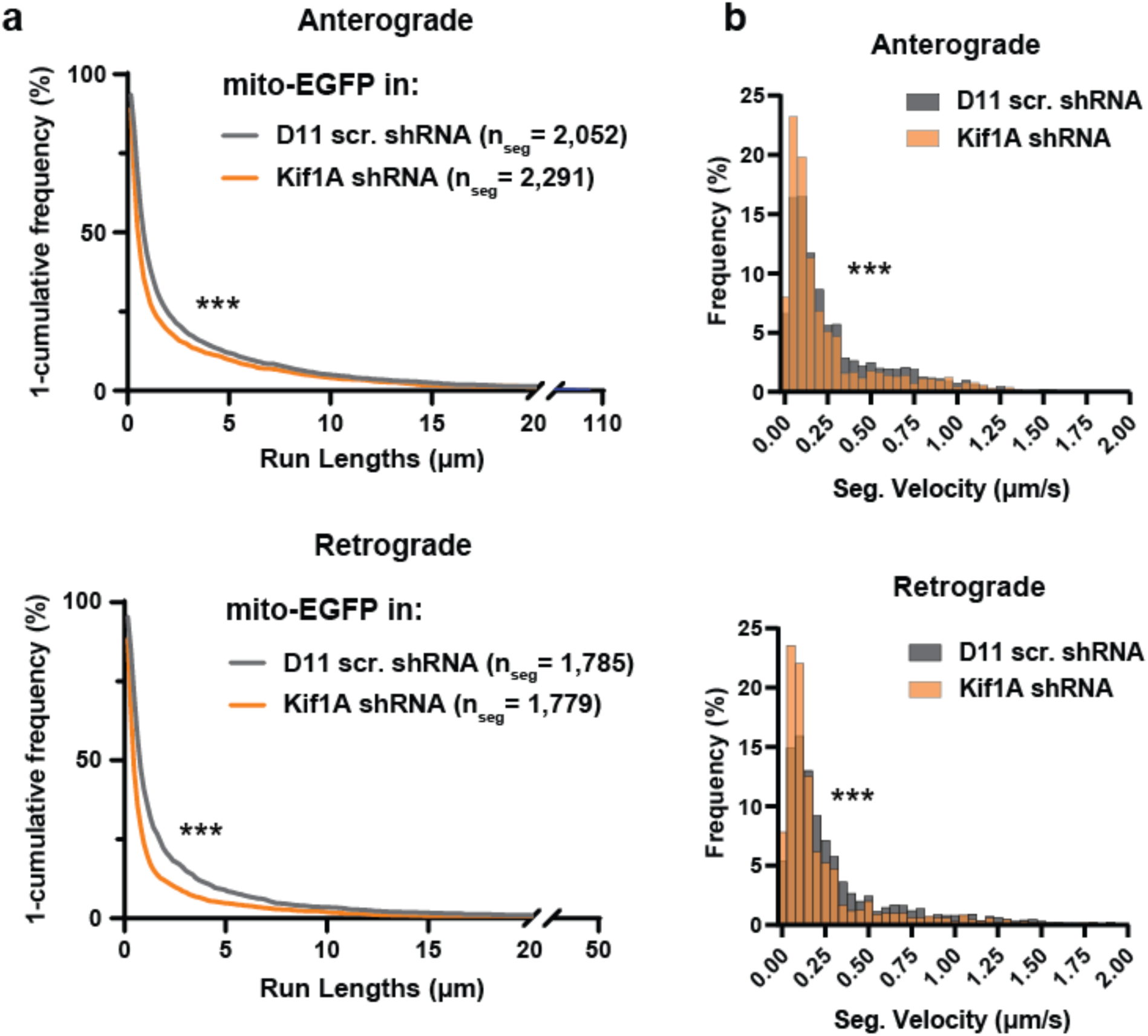
Mitochondrial transport in mammalian axons requires KIF1A. **a** Anterograde (top) and retrograde (bottom) segmental run lengths of mito-EGFP in D11 scrambled (scr.) shRNA vs Kif1A shRNA, plotted as 1-cumulative frequency (%). **b** Anterograde (top) and retrograde (bottom) segmental velocity of mito-EGFP in D11 scrambled (scr.) shRNA vs Kif1A shRNA plotted as relative frequency histogram. ***p<0.001, Kolmogorov-Smirnov (K-S) test.

**Supplementary Fig. S3:**
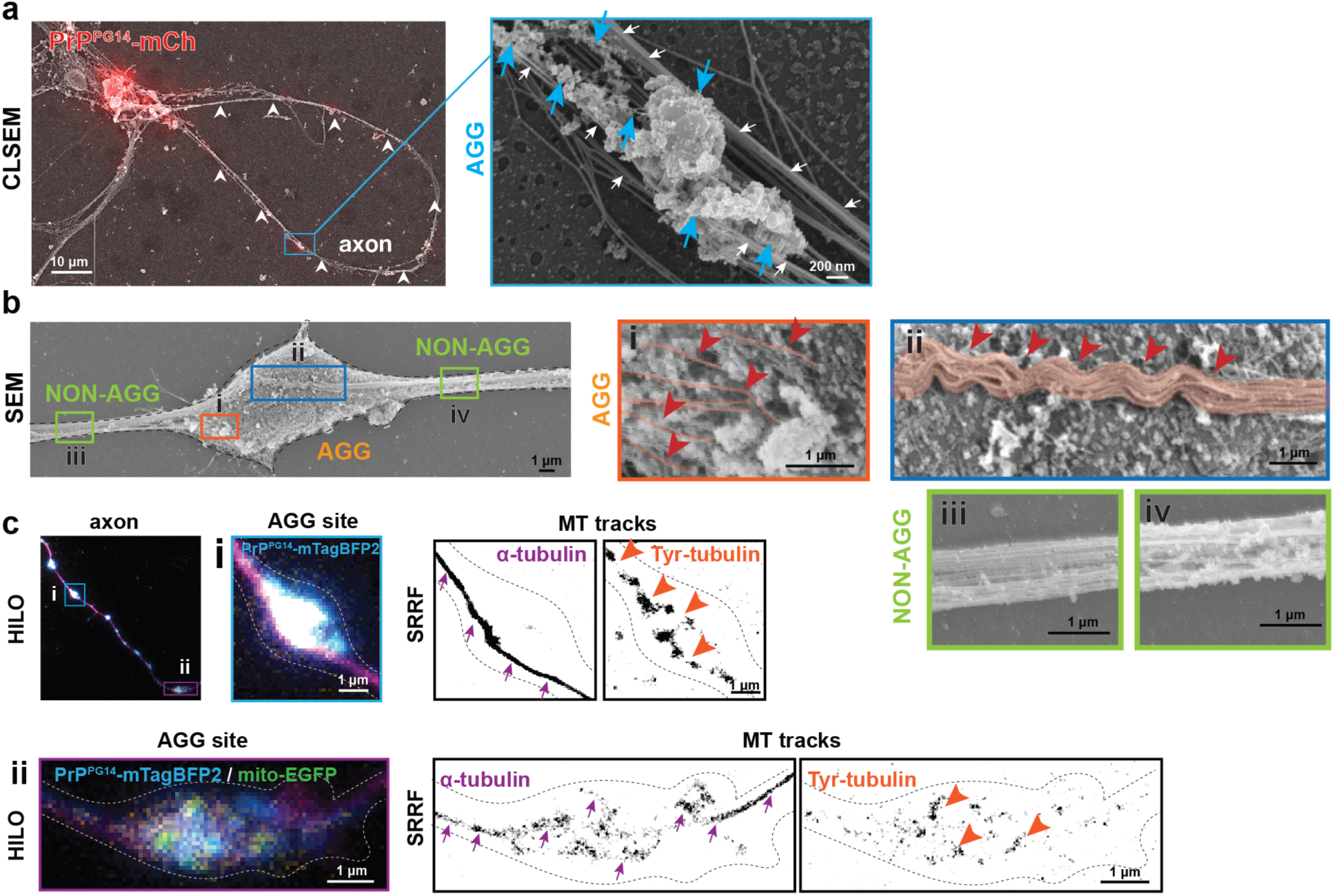
Ultrastructure of the axonal cytoskeleton at PrP^PG14^ aggregate sites. **a** Membrane-extracted CLSEM image of a neuron transfected with PrP^PG14^-mCh (red). White arrowheads indicate axons. Blue inset (right) shows an AGG site. White arrows point to MT tracks. Blue arrows point to actin surrounding PrP^PG14^ aggregates. **b** Membrane-extracted SEM image of an axonal region of a neuron transfected with PrP^PG14^-mCh. Insets (i-ii) show AGG sites. Red arrowheads and pseudo-color highlights show broken (i) or bending (ii) MT tracks. Insets (iii-iv) show NON-AGG sites. **c** HILO image of an axon from a neuron co-transfected with PrP^PG14^-mTagBFP2 and mito-EGFP, co-immunostained with alpha tubulin and antibodies against tyrosinated (Tyr) tubulin. Inset (i) shows AGG site and SRRF processed images of alpha tubulin and Tyr-tubulin channels. Inset (ii) shows a HILO image of PrP^PG14^-mTagBFP2 and mito-EGFP at AGG site and SRRF processed images of alpha tubulin and Tyr-tubulin channels. Arrows point to straight/intact MT tracks. Arrowheads point to broken MT tracks.

**Supplementary Fig. S4:**
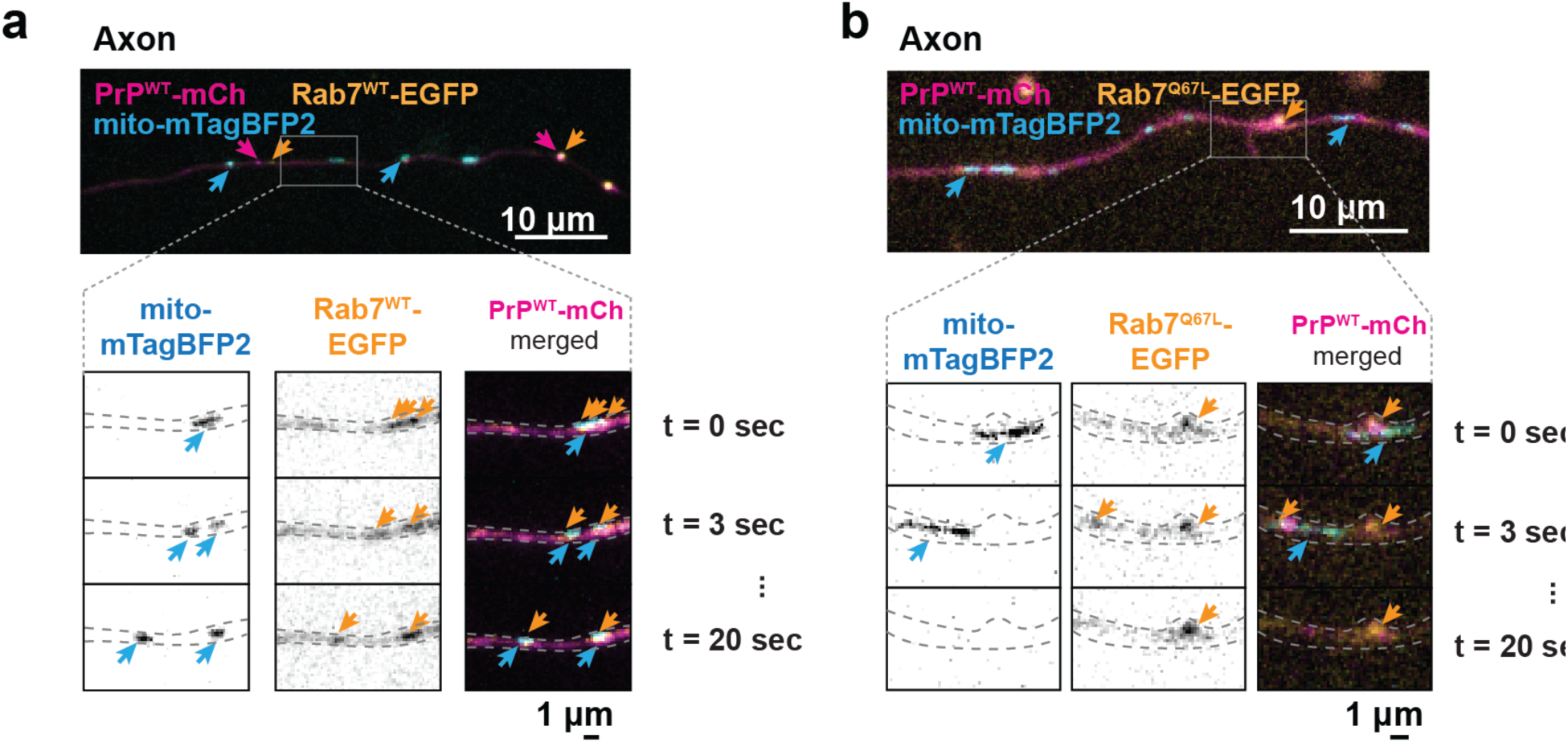
Mitochondrial fission in PrP^WT^ axons. **a** First-frame images of axons co-expressing PrP^WT^-mCh, Rab7^WT^-EGFP, and mito-mTagBFP2 and representative frames from time-lapse videos at WT or AGG sites (insets). **b** First-frame images of axons co-expressing PrP^WT^-mCh, Rab7^Q67L^-EGFP, and mito-mTagBFP2 and representative frames from time-lapse videos at WT or AGG sites (insets). Arrows point to corresponding organelles in the first-frame images and time-lapse insets.

